# Target- and network-specific downstream effects of personalized precision transcranial magnetic stimulation: Evidence from concurrent functional neuroimaging

**DOI:** 10.64898/2026.11.10.765235

**Authors:** Maximilian Lueckel, Angela Radetz, Umair Hassan, Kenneth Yuen, Florian Müller-Dahlhaus, Raffael Kalisch, Til Ole Bergmann

## Abstract

**Background:** Personalized targeting of transcranial magnetic stimulation (TMS) based on functional connectivity (FC) to the subgenual anterior cingulate cortex (sgACC) is promising for depression treatment, but direct evidence that precisely engaging an individually defined FC target produces specific downstream modulation is lacking.

**Objective:** To test whether stimulating individually defined cortical targets with strong negative or positive FC to sgACC/ventromedial prefrontal cortex (vmPFC) produces target- and network-specific downstream BOLD effects compared to nearby control targets.

**Methods:** Twenty healthy participants underwent concurrent TMS-fMRI. For each participant, we identified a dorsolateral PFC target with strong negative FC and a dorsomedial PFC target with strong positive FC to sgACC/vmPFC as well as control targets with weaker sgACC/vmPFC-FC strength, but matched peripheral co-stimulation experience. Coil positions and orientations were optimized using electric field simulations. Single TMS pulses were applied during concurrent fMRI, and BOLD responses were compared between conditions.

**Results:** Stimulation of negative FC targets produced significantly stronger negative BOLD modulation in sgACC/vmPFC and dorsomedial PFC compared to control targets. Positive FC targets showed no significant differences from controls. The spatial pattern of negative FC target-related BOLD effects significantly resembled a previously described convergent depression circuit, while positive FC targets produced an anticorrelated pattern.

**Conclusions:** Personalized precision TMS targeting produces target- and network-specific downstream effects, supporting the mechanistic premise of FC-guided TMS.

**Highlights:**

- Negative FC targets (vs. controls) evoked negative sgACC/vmPFC-BOLD modulations.
- Positive FC targets (vs. controls) showed no target-specific downstream modulation.
- TMS-evoked BOLD patterns were consistent with anticorrelated network architectures.
- Control targets accounted for peripheral co-stimulation confounds.
- Coil placements were individually optimized using FC and E-field simulations.

## 1 Introduction

Transcranial magnetic stimulation (TMS) is a non-invasive brain stimulation technique with established therapeutic efficacy in several psychiatric and neurological disorders (1). For example, high-frequency repetitive TMS (rTMS) targeting the left dorsolateral prefrontal cortex (DLPFC) is an evidence-based treatment for treatment-resistant depression (1,2). However, clinical response rates remain modest and variable (3–5). A key factor likely contributing to this variability is the spatial placement of the TMS coil, which determines not only the specific cortical site but also the connected functional network(s) engaged in each individual (6–9).

Over the past decade, this insight has motivated a shift towards personalized TMS targeting (9). Rather than relying on heuristic scalp measurements or group average coordinates, stimulation sites are increasingly defined based on each individual’s resting-state functional connectivity (FC) to disorder-relevant brain regions or circuits, to, for example, engage the subgenual anterior cingulate cortex (sgACC) in depression (10–12). Such individual connectivity-guided approaches have shown promising clinical efficacy (13,14), superior to conventional scalp-based targeting (15,16). In addition, electric field (E-field) modelling is increasingly used to refine coil position and orientation and maximize engagement of target sites and networks (6,17–19).

This personalized precision targeting approach rests on the mechanistic premise that precisely stimulating a person-specific cortical volume, rather than a general cortical region (e.g., DLPFC), produces qualitatively different and therapeutically more effective modulation of the connected downstream target (e.g., sgACC) and/or network. However, so far there is only indirect evidence, primarily from (retrospective) associations between proximity of individual stimulation sites to FC-defined coordinates and clinical improvement (10–12,20,21) as well as from studies showing that TMS modulates connected deep regions in general (e.g., 22–26). To date, no study has directly tested whether precisely engaging an individually defined FC target produces downstream network modulation beyond what would result from stimulating an adjacent non-target site within the same cortical region.

Furthermore, it remains unclear whether the direction (activation/deactivation) and magnitude of downstream effects depend on the specific FC profile of the cortical network node being primarily engaged. Retrospective work has linked stronger negative FC between the stimulation site and sgACC to better antidepressant response, establishing negative sgACC-FC as the dominant target selection criterion in current FC-guided protocols (10,11,27). Yet, recent concurrent TMS-fMRI work found stimulation sites with positive rather than negative FC to sgACC to produce stronger evoked sgACC responses in depressed patients (28,29), suggesting that positively connected cortical targets may influence downstream regions as well (30). Whether negative and positive FC cortical targets differ regarding their downstream effects on a given deep brain target and whether a specific connectivity profile should be preferably selected yet needs to be determined.

Here, we combined individualized, FC-guided target identification and E-field-optimized precision targeting with concurrent TMS-fMRI to causally probe FC target specificity in downstream brain modulation (31,26). For each participant, we identified two individually defined cortical targets based on their connectivity to sgACC and adjacent ventromedial prefrontal cortex (vmPFC): (i) a dorsolateral PFC site with strong negative FC, and (ii) a dorsomedial PFC site with strong positive FC. For each FC target, we additionally defined a nearby control site, with matched peripheral sensory experience but weaker sgACC/vmPFC connectivity. This design allowed us to test (i) whether precisely stimulating the FC-defined site produces distinct downstream modulation compared to its immediate cortical surroundings, and (ii) whether this effect depends on the functional connectivity profile (negative vs. positive) of the targeted network.

## 2 Methods

### 2.1 Sample

Twenty healthy participants (15 female; age: M = 25.35 years, SD = 3.34 years, range: 20-32 years) with no history of psychiatric or neurological disorders were recruited. A physician screened all participants for MRI and TMS eligibility (see **Supplementary Methods** for inclusion and exclusion criteria). Study procedures were approved by the local ethics committee in accordance with the Declaration of Helsinki (ethics vote no. 2019-14452), and all participants gave written informed consent before study enrollment.

### 2.2 Experimental Procedure

#### Intake session

Before TMS-fMRI, participants underwent resting-state functional and anatomical MRI (see **Supplementary Methods** for sequence parameters, data processing, and detailed procedures).

#### Concurrent TMS-fMRI sessions

Participants underwent two TMS-fMRI sessions, each comprising two runs. Each session targeted either the left *dorsolateral* PFC target with strong *negative* FC to sgACC/vmPFC and its control target, or the left *dorsomedial* PFC target with strong *positive* FC to sgACC/vmPFC and its control target (see below for target selection). The order of FC and control target stimulation was counterbalanced within sessions, and positive/negative FC target order was counterbalanced across sessions, which were separated by at least 2 days.

Stimulation intensity was individually adjusted for FC targets based on E-field simulations and tolerability. For control targets, intensity was set to match the subjective experience (unpleasantness, loudness) of FC target stimulation as closely as possible. In each run, we applied 65 single biphasic TMS pulses, with a jittered inter-pulse interval of 10-15 s. Each suprathreshold TMS pulse was preceded by two low-intensity, sub-threshold pulses (∼35% of target intensity) to reduce startle-related head movements and eye blinks (**Figure S1; Table S1**). Immediately after each run, participants rated (i) stimulation unpleasantness, (ii) TMS click loudness, (iii) TMS-related muscle twitches, and (iv) tiredness (0–10). See **Supplementary Methods** for detailed procedures.

### 2.3 TMS Targeting Optimization

#### Functional connectivity analysis

Individual FC maps for target identification and coil placement optimization were computed from participants’ preprocessed intake MRI data. A normative, whole-brain sgACC-/vmPFC-based FC map derived from a large normative sample using previously reported sgACC (10) and vmPFC (32) ROIs (**Figure S2A**) was used as a weighted seed in each participant’s surface-projected resting-state fMRI data (**Figure 1A-B, Figure S2B-C**) (33,21,34). See **Supplementary Methods** for details.

**Figure 1.**
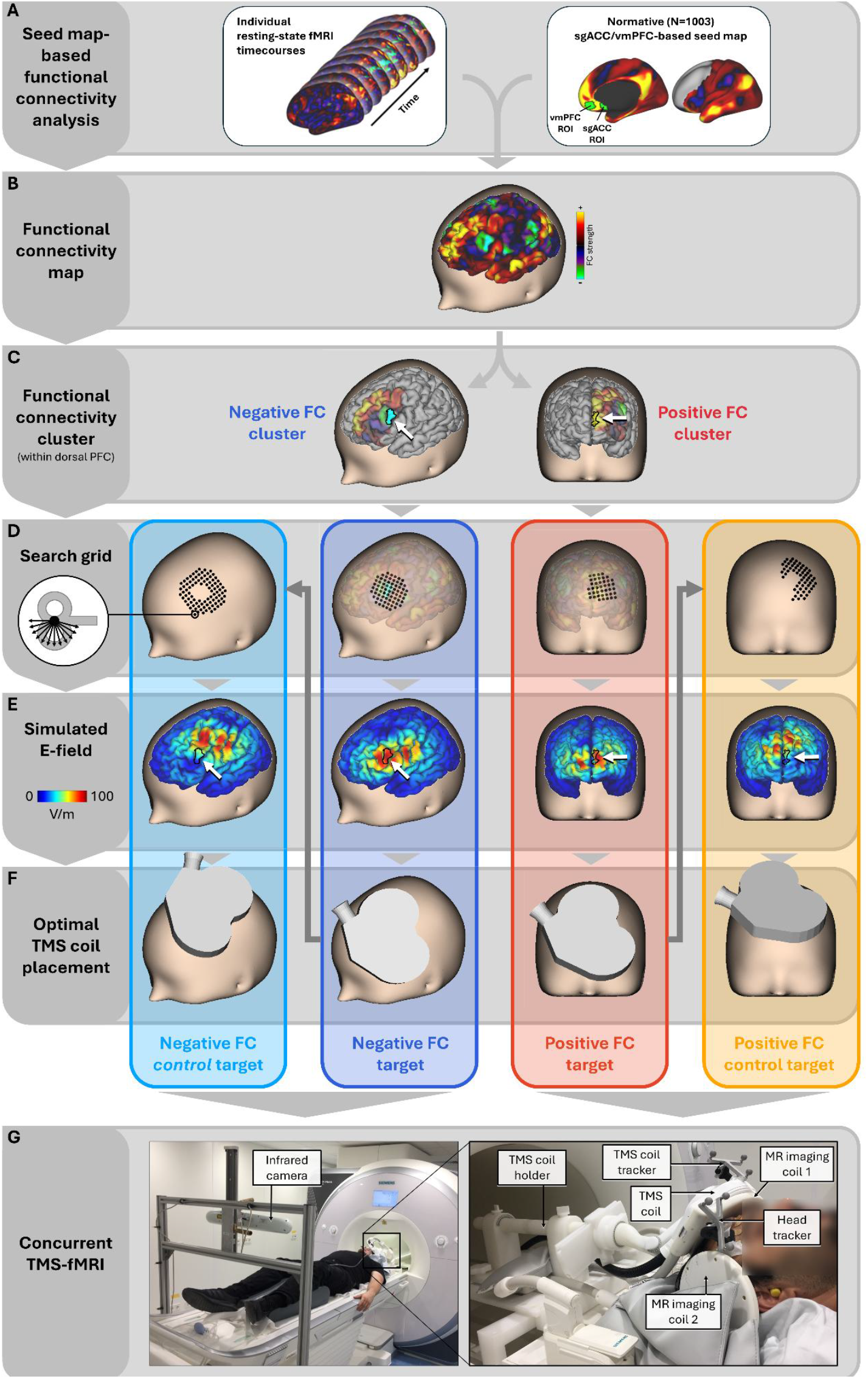
Overview of the methodological approach. **A**: Functional connectivity analysis using individual resting-state fMRI data and a normative sgACC-/vmPFC-based FC map (excluding the dorsal PFC) as seed. **B**: Resulting individual FC map (from a representative participant). **C**: Individually identified largest clusters of negative and positive FC with the sgACC/vmPFC (seed map) within dorsal PFC. **D**: Condition-specific search grids of potential TMS coil positions and orientations, for which E-field simulations were run. **E**: Simulated E-fields for the condition-specific optimal combinations of TMS coil position and orientation. **F**: Condition-specific, individually optimized TMS coil placements. **G**: Concurrent TMS-fMRI setup (photographs depict members of the author team).

#### TMS target identification

Individual FC maps were first multiplied with individual sulcal depth maps, upweighting FC clusters on the gyral crown (**Figure S3**) (cf. 6), where the TMS-induced E-field is maximal (35,36). Within the left dorsal PFC (**Figure 1C**), we identified the largest clusters of negative and positive sgACC/vmPFC-FC. See **Supplementary Methods** for details.

#### Electric field modelling

Using realistic individualized head models in SimNIBS (version 4.0) (37), E-field simulations were run across numerous potential coil positions and orientations surrounding each FC cluster. After identifying the optimal FC target placement (see below), a second set of simulations searched surrounding positions and orientations for the corresponding control placement (see below; **Figure 1D**). See **Supplementary Methods** for details.

#### Selection of optimal TMS coil placements

The optimal coil placement for each FC cluster maximized both E-field magnitude within the target cluster and overlap between the whole-brain sgACC/vmPFC-FC map and the E-field map (**Figure 1E**). Unlike targeting procedures that treat the cortical target as a single coordinate or spatially restricted ROI, this approach additionally accounts for the whole-brain topography of both FC and induced E-field, maximizing effects on target areas while largely sparing off-target areas (**Figure 1E**).

Control placements were determined similarly, maximizing overlap between an inverted FC map and the E-field map. We selected the coil position/orientation that would (i) stimulate PFC with, on average, weaker sgACC/vmPFC-FC than the corresponding target cluster; (ii) lie as close as possible to the FC-optimized placement to match peripheral co-stimulation; and (iii) induce an E-field as weak as possible within the corresponding FC target cluster to avoid effectively co-stimulating it. This approach aimed to simultaneously maximize experimental variance and minimize confounding variance between conditions. See **Supplementary Methods** for details.

#### Determination of stimulation intensity

Stimulation intensity was individually determined from the same E-field simulations, aiming for a mean E-field strength of 100 V/m within the FC target clusters (38–40,6). If the simulation-derived intensity exceeded technical limits or tolerability, we used the highest technically possible or tolerable intensity. See **Supplementary Methods** for details.

### 2.4 Concurrent TMS-fMRI

#### Acquisition, preprocessing, and first-level analysis

TMS-fMRI was acquired using two TMS-compatible 7-channel RF surface arrays (**Figure 1G**) (41) with a multi-band, multi-echo EPI sequence, and an MRi-B91 TMS coil (MagVenture, Denmark). An MR-compatible neuronavigation system (Localite GmbH, Germany) enabled precise coil navigation to the individually optimized placements (**Figure 1G**). The TMS device (MagPro X100, MagVenture, Denmark) was controlled and synchronized with MRI acquisition using the BEST toolbox (42). Data preprocessing included NORDIC denoising (43), bias field correction, fMRIPrep preprocessing (44), and tedana (45). At the first level, TMS pulses were modelled as instantaneous events (i.e., delta functions) convolved with a canonical hemodynamic response function, yielding one contrast map per stimulation condition and participant. See **Supplementary Methods** for sequence parameters as well as procedural and processing details.

#### Second-level analysis

Condition-specific first-level contrast maps were entered into second-level GLMs. One-sample t-tests assessed effects of each stimulation condition separately, and paired t-tests contrasted FC target against respective control target stimulation. Differences between negative and positive FC targets were tested using the interaction contrast: [(negative FC target vs. negative FC *control* target) vs. (positive FC target vs. positive FC *control* target)]. Each model included subject- and condition-specific unpleasantness and loudness ratings as covariates of no interest. Multiple comparisons were corrected using cluster-level family-wise error (FWE) correction (cluster-forming threshold p < .001, uncorrected) (46).

#### Spatial similarity

To assess the potential clinical relevance of the resulting contrast maps, spatial correlation (r) between each unthresholded group-level t-map and a previously reported convergent depression circuit (47) was tested for significance using spin tests (10,000 rotations) (48). Two-sided *p*_spin_-values were Bonferroni-corrected across maps.

### 2.5 Statistical Analysis

Derived measures (e.g., E-field values, behavioral ratings, beta values from contrast maps, etc.) were analyzed in R (version 4.4.3) (49). Conditions were compared using paired t-tests if the paired differences were normally distributed (Shapiro-Wilk *p* > .05) and Wilcoxon signed-rank tests otherwise. P-values were Bonferroni-corrected within each set of comparisons and are reported as corrected values (p_Bonf_; α = .05). Effect sizes are reported as Cohen’s d for repeated measures (50) for t-tests and as matched-pairs rank-biserial correlations for Wilcoxon tests, each with a Bonferroni-adjusted confidence interval. Target type (FC target vs. control target) × FC valence (positive vs. negative) interactions were tested using two-way repeated-measures ANOVAs with covariates (loudness and unpleasantness ratings), with partial η² (η²_p_) as effect size measure.

## 3 Results

### 3.1 TMS targets and coil placements

**Figure 2A** illustrates the interindividual variability in center locations of the largest negative (blue) and positive (red) FC clusters within the left dorsal PFC. The personalization procedure resulted in considerable interindividual variability in both TMS coil position and orientation (**Figure 2B**; see **Supplementary Results** and **Figure S5** for information on positional and rotational variability).

**Figure 2.**
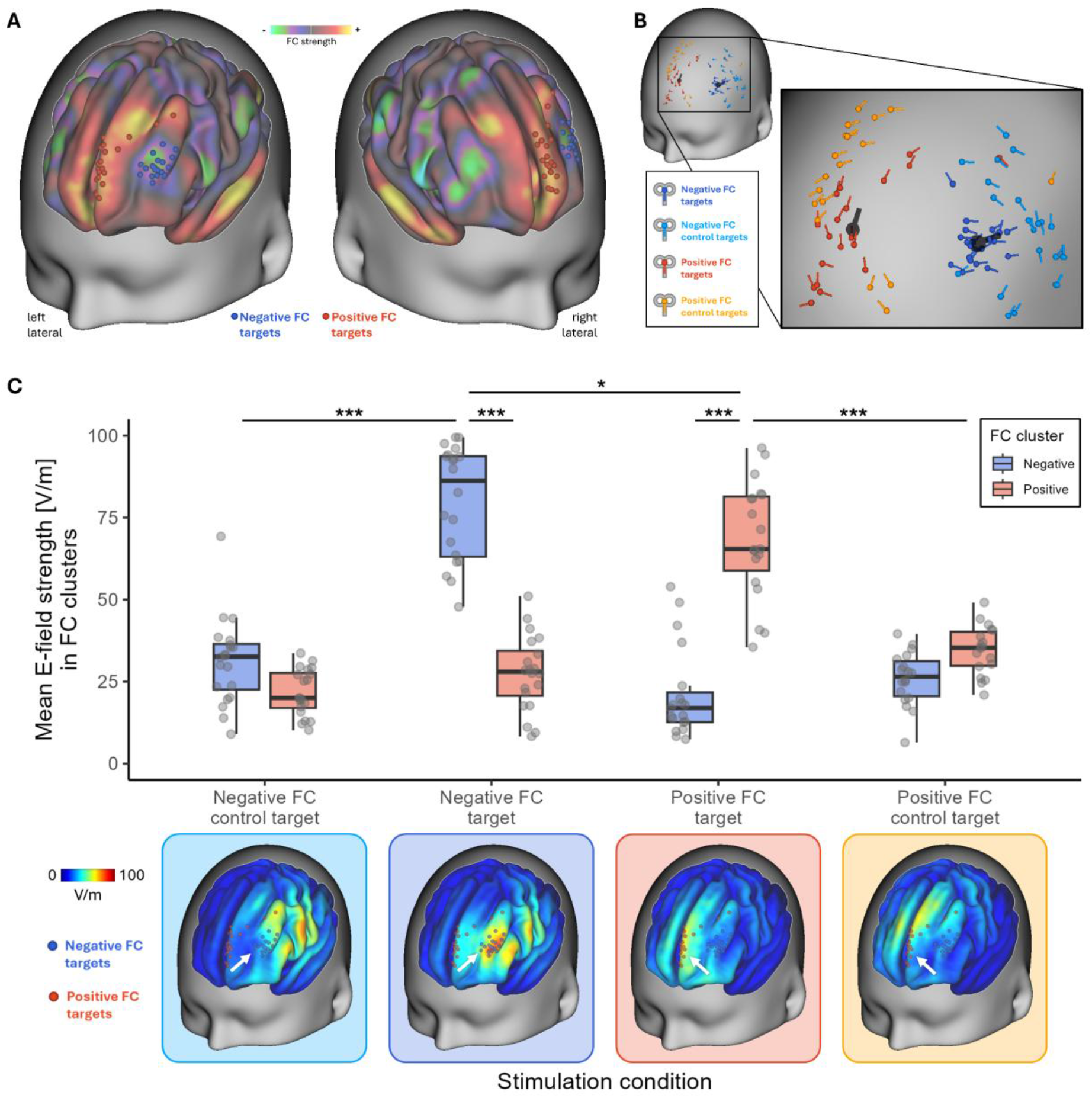
Overview of TMS targets, coil placements, and average simulated TMS-induced electric fields per stimulation condition for the whole sample of N=20 participants. **A**: Blue and red dots represent the geometric centers of the individually identified negative and positive FC target clusters, respectively, overlaid on the sample-averaged, sgACC/vmPFC seed map-based FC map. **B**: Schematic illustration of the condition-specific, individually optimized TMS coil placements (position and orientation), with the larger black coil representations indicating the average coil placement for the respective negative and positive FC target conditions. See Supplementary Results and Figure S5 for information on positional and rotational deviations. **C:** Boxplots of mean E-field values in the individually identified negative (blue) and positive (red) FC clusters (*n* = 20 for negative FC conditions, *n* = 19 for the positive FC target, *n* = 18 for the positive FC control target; top) and sample-averaged simulated E-fields visualized on the cortical surface (bottom). White arrows indicate clusters of the geometric centers of negative and positive FC target clusters. Brackets indicate paired t-tests, Bonferroni-corrected for five comparisons (see Results section 3.2). \*\*\**p*_Bonf_ < .001. \**p*_Bonf_ < .05.

### 3.2 TMS-induced electric fields

**Figure 2C** shows mean E-field values within the individual negative and positive FC clusters and sample-averaged simulated E-fields per condition (see **Table 1** for means and SDs). Optimized coil placements induced strong E-fields within their respective target clusters, whereas control coil placements produced significantly weaker E-fields within the negative (mean difference = 48.5 V/m [39.5, 57.5], *t*(19) = 15.46, *p*_Bonf_ < .001, *d* = 3.08 [1.79, 4.38]) and positive (mean difference = 34.5 V/m [22.8, 46.2], *t*(17) = 8.52, *p*_Bonf_ < .001, *d* = 2.33 [0.89, 3.77]) FC clusters, respectively. Likewise, coil placements optimized for one FC cluster induced significantly weaker E-fields in the FC cluster of opposite valence (negative FC target stimulation: mean difference = 52.3 V/m [43.1, 61.5], *t*(19) = 16.24, *p*_Bonf_ < .001, *d* = 3.41 [1.92, 4.90]; positive FC target stimulation: mean difference = 47.2 V/m [35.6, 58.9], *t*(18) = 11.64, *p*_Bonf_ < .001, *d* = 2.92 [1.35, 4.49]). This shows that the optimization procedure maximized E-field strength within target clusters while minimizing it in non-target clusters, as intended. Since the procedure did not explicitly match E-fields across the two FC target clusters, E-fields in negative FC clusters during negative FC target stimulation happened to be significantly stronger than those in the positive FC clusters during positive FC target stimulation (mean difference = 10.6 V/m [0.1, 21.2], *t*(18) = 2.90, *p*_Bonf_ = .047, *d* = 0.61).

**Table 1.** Descriptive statistics (mean and standard deviation) of mean E-field strength in the individual FC clusters, stimulation intensity and subjective ratings, and mean beta estimates in regions of interest and significant clusters, for each stimulation condition.

|  | Stimulation condition |  |  |  |  |  |  |  |
|---|---|---|---|---|---|---|---|---|
|  | Negative FC<br>control target<br>(n = 20) |  | Negative FC<br>target<br>(n = 20) |  | Positive FC<br>target<br>(n = 19) |  | Positive FC<br>control target<br>(n = 18) |  |
|  | M | SD | M | SD | M | SD | M | SD |
| **Mean E-field [V/m] values in FC clusters (cf. Figure 2)** |  |  |  |  |  |  |  |  |
| Negative FC cluster | 31.31 | 13.22 | 79.83 | 17.22 | 21.10 | 13.91 | 25.88 | 8.16 |
| Positive FC cluster | 21.80 | 7.07 | 27.50 | 11.49 | 68.34 | 17.86 | 34.58 | 7.51 |
| **Confound variables (cf. Figure 3)** |  |  |  |  |  |  |  |  |
| % MSO [0-100] | 82.45 | 12.76 | 84.55 | 14.07 | 88.58 | 13.85 | 88.50 | 16.27 |
| Unpleasantness [0-10] | 4.30 | 2.27 | 4.43 | 2.03 | 4.03 | 2.31 | 3.67 | 2.57 |
| Loudness of the TMS click sound [0-10] | 4.70 | 1.58 | 4.73 | 1.52 | 4.63 | 1.50 | 4.00 | 1.53 |
| **Mean beta estimates in regions of interest (cf. Figure 4)** |  |  |  |  |  |  |  |  |
| sgACC ROI | 0.15 | 0.94 | -0.88 | 1.22 | -0.30 | 0.75 | -0.44 | 0.46 |
| vmPFC ROI | -0.13 | 1.55 | -1.98 | 2.29 | -0.76 | 1.64 | -0.39 | 1.15 |
| Significant sgACC/vmPFC cluster | 0.67 | 1.27 | -1.29 | 1.10 | -0.10 | 0.95 | -0.72 | 0.81 |
| Significant dmPFC cluster | -1.06 | 1.52 | -3.58 | 3.05 | -0.66 | 0.91 | -1.20 | 1.14 |
| Significant bilateral parietal cortex cluster | -3.06 | 2.01 | -0.49 | 1.98 | -1.69 | 0.87 | -0.55 | 0.95 |

*Note.* Means and SDs are based on all available data per condition, while statistical tests only used complete pairs.

### 3.3 Confounds

As illustrated in **Figure 3** (see also **Table 1**), realized stimulation intensity (% MSO), unpleasantness, and perceived TMS click loudness did not differ significantly between negative FC targets and controls, positive FC targets and controls, and negative and positive FC targets (all *p*_Bonf_ ≥ .078; see **Supplementary Results** for pairwise comparison results). Together, these results provide no evidence for systematic differences in unspecific peripheral co-stimulation between conditions. Nevertheless, unpleasantness and loudness ratings were included as covariates of no interest in all group-level analyses.

**Figure 3.**
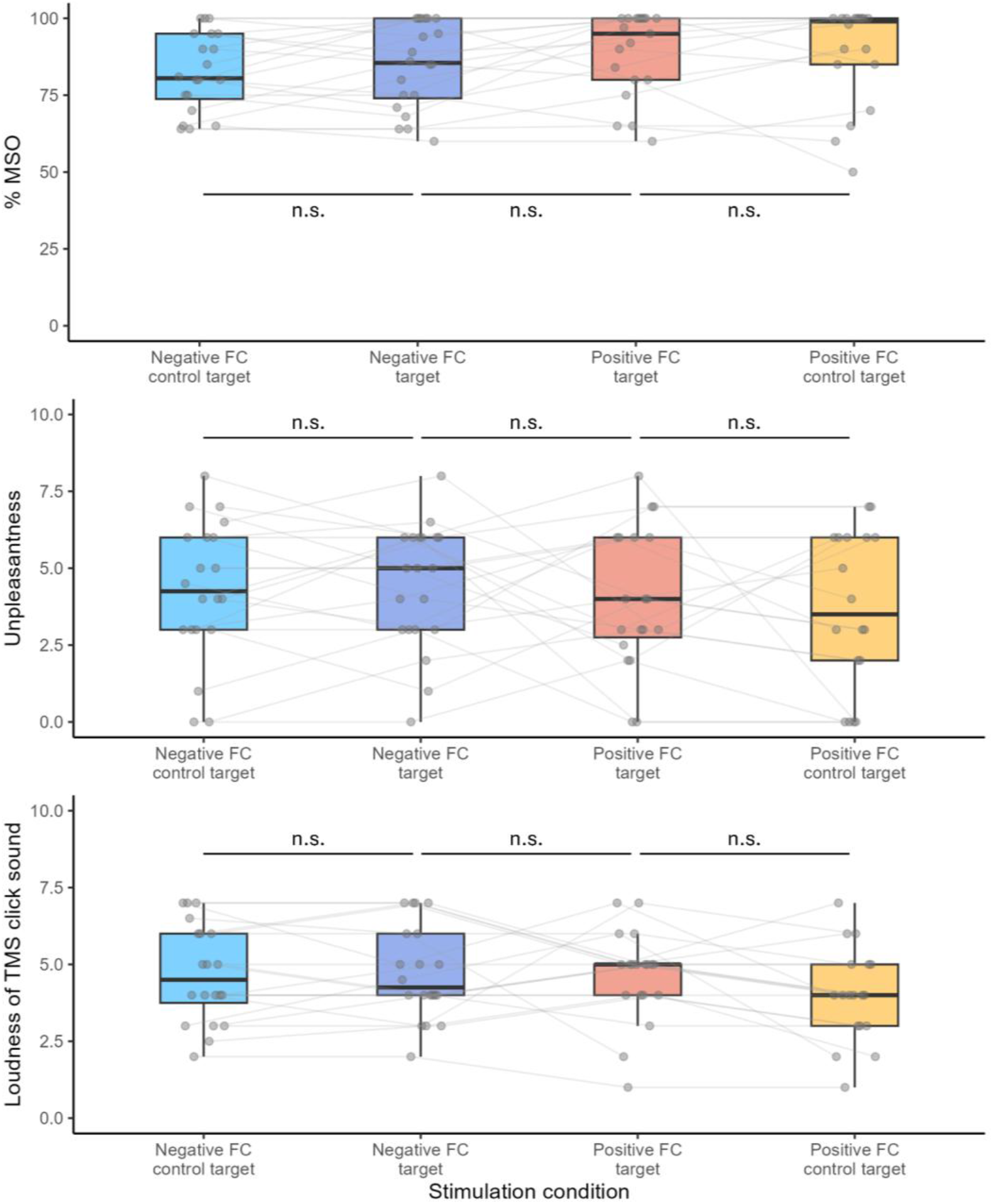
Boxplots of confound measures for each stimulation condition. Top: Stimulation intensity (% maximum stimulator output, MSO; 0-100). Middle: Subjective ratings of stimulation unpleasantness (0–10). Bottom: Subjective ratings of the loudness of the TMS click sound (0–10). Grey dots represent individual participants, and lines connect values from the same participant (*n* = 20 for negative FC conditions, *n* = 19 for the positive FC target, *n* = 18 for the positive FC control target). Brackets indicate paired t-tests or, for non-normally distributed differences, Wilcoxon signed-rank tests, Bonferroni-corrected for three comparisons per measure (see Supplementary Results for pairwise comparison results). n.s. = not significant (*p*_Bonf_ > .05).

### 3.4 Concurrent TMS-fMRI

#### Condition-specific results

Across all four stimulation conditions, we consistently observed significant TMS-related positive BOLD modulations in the auditory network (primary/secondary auditory cortex, i.e., Heschl’s gyrus, planum temporale, superior temporal gyrus) and the cingulo-opercular network (CON; anterior/posterior insula, dorsal anterior cingulate cortex (dACC), supramarginal gyrus), as well as subcortically in the bilateral thalamus, midbrain, brainstem, and cerebellum. Negative BOLD modulations were consistently found in the somatomotor network (face, hand, and foot representations in pre- and postcentral gyri, central sulcus, paracentral lobule) and, less consistently, but most pronounced for the negative FC target condition, within the default mode network (DMN), including temporal and dorso-/ventromedial prefrontal regions encompassing sgACC and vmPFC (**Figure 4A**). For peak coordinates of significant clusters, see **Supplementary Tables S2-S9**.

**Figure 4.**
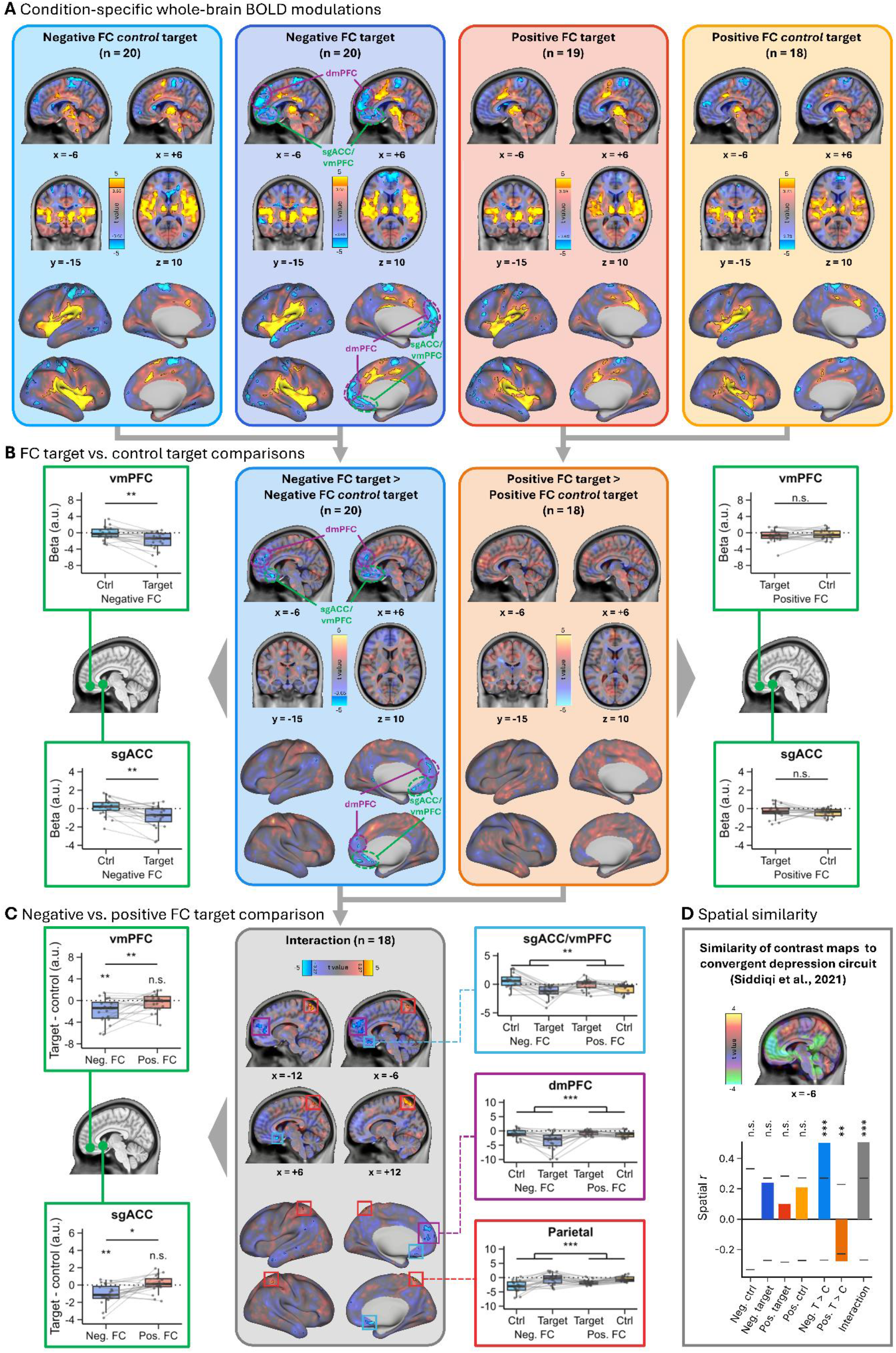
TMS-fMRI BOLD modulation results. **A**: Whole-brain BOLD modulations separately for each stimulation condition (one-sample t-tests; negative FC control target and negative FC target: n = 20; positive FC target: n = 19; positive FC control target: n = 18), shown on sagittal (x = −6, +6), coronal (y = −15) and axial (z = 10) slices of the MNI template and on the inflated cortical surface. **B**: Whole-brain comparison of FC target vs. control target stimulation (paired t-tests), separately for negative (n = 20) and positive (n = 18) FC targets, and beta values extracted from the a priori vmPFC (32) and sgACC (10) ROIs (green frames; ROI locations indicated on the sagittal slice). ROI comparisons were adjusted for unpleasantness and loudness ratings and Bonferroni-corrected for two comparisons per ROI. **C**: Whole-brain comparison of negative vs. positive FC targets, each relative to their control targets [(negative FC target > negative FC control target) > (positive FC target > positive FC control target); n = 18]. Colored boxes mark the three significant clusters: sgACC/vmPFC (light blue), dmPFC (purple) and parietal cortex (red). Left: Beta differences between FC target and control target stimulation in the a priori vmPFC and sgACC ROIs, separately for negative (n = 20) and positive (n = 18) FC targets. Stars above boxes indicate FC target vs. control comparisons (Bonferroni-corrected for two comparisons per ROI); brackets indicate the target type × FC valence interaction (n = 18; Bonferroni-corrected for two ROIs). All ROI tests were adjusted for unpleasantness and loudness ratings. Right: Beta values extracted from the three significant interaction clusters (n = 18). Brackets indicate the significance of the whole-brain interaction cluster (cluster-level *p*_FWE_). Note that, as the clusters were selected based on this contrast, no further statistical tests were performed on the extracted values. **D**: Top: Convergent depression circuit (47) shown on a sagittal slice (x = −6) of the MNI template. Bottom: Spatial correlation between each unthresholded group t-map and the convergent depression circuit. Bars show observed spatial *r* and short horizontal lines indicate the Bonferroni-corrected significance thresholds derived from 10,000 spatial rotations of the circuit map (spin test; two-sided, α = .05/7). Stars mark Bonferroni-corrected spin-test *p*-values. Neg./Pos. = negative/positive FC; ctrl = control target; T > C = FC target > control target. For transparency, significant BOLD modulation clusters (cluster-level *p*_FWE_ < .05, cluster-forming threshold *p* < .001, uncorrected) are shown with black outlines in front of the translucent unthresholded maps (83) throughout the figure. For the surface illustrations, the same voxel-wise threshold as for the volume representations was applied without removing clusters below the cluster-extent threshold. Small clusters may therefore appear on the surface but should not be interpreted. In the boxplots, grey dots represent individual participants and lines connect values from the same participant. Dotted lines mark zero. All ROI and cluster beta values are plotted unadjusted. \**p* < .05, \*\**p* < .01, \*\*\**p* < .001 (Bonferroni-corrected or cluster-level FWE-corrected, as specified above); n.s. = not significant. Abbreviations: dmPFC = dorsomedial prefrontal cortex; sgACC = subgenual anterior cingulate cortex; vmPFC = ventromedial prefrontal cortex.

#### FC target vs. control target comparisons

Stimulation of negative FC targets produced significantly stronger negative BOLD modulations than stimulation of their control targets in dorso- and ventromedial prefrontal areas, including sgACC and vmPFC. For peak coordinates, see **Supplementary Table S10**. Beta estimates from the a priori sgACC and vmPFC ROIs (**Table 1**), adjusted for differences in unpleasantness and loudness ratings, confirmed significantly more negative values for negative FC targets than for their control targets (sgACC: adjusted difference = −1.03 [−1.69, −0.36], *t*(17) = −3.78, *p*_Bonf_ = .003; vmPFC: adjusted difference = −1.80 [−3.00, −0.59], *t*(17) = −3.66, *p*_Bonf_ = .004; **Figure 4B**, left). These effects were virtually unchanged by the adjustment (unadjusted differences: −1.03 and −1.86), and rating differences did not predict them (all *p* ≥ .147).

In contrast, TMS-induced BOLD modulations for positive FC targets did not differ significantly from their control targets anywhere in the brain, including the sgACC (adjusted difference = 0.26 [−0.45, 0.98], *t*(15) = 0.91, *p*_Bonf_ = .752) or vmPFC (−0.16 [−1.23, 0.91], *t*(15) = −0.38, *p*_Bonf_ > .99) ROIs (**Figure 4B**, right; **Table 1**)

#### Negative vs. positive FC target comparison

Comparing negative and positive FC target stimulation (both relative to their controls) revealed significantly stronger negative BOLD modulations in dorsomedial as well as ventromedial PFC and subgenual ACC for negative FC targets. This interaction was driven by more negative beta values for negative FC targets than for their control targets, with no such clear difference between positive FC targets and their control targets (cf. above; **Figure 4C**, right; **Table 1**). Importantly, this interaction was also significant in the independently defined a priori ROIs (sgACC: adjusted interaction contrast = −1.41 [−2.54, −0.27], t(15) = −3.09, *p*_Bonf_ = .015, η²_p_ = .39; vmPFC: −2.18 [−3.81, −0.55], t(15) = −3.33, *p*_Bonf_ = .009, η²_p_ = .43; n = 18; **Figure 4C**, left)

We also found significantly stronger positive BOLD modulations in bilateral parietal cortex for negative FC targets than for positive FC targets (each relative to their controls). For this interaction effect, the pattern was reversed: negative FC control targets produced more negative beta values than negative FC targets, whereas positive FC targets produced more negative values than their controls (**Figure 4C**, right; **Table 1**). For peak coordinates of significant interaction clusters, see **Supplementary Tables S11-S12**.

#### Spatial similarity with convergent depression circuit

Both the BOLD modulation pattern evoked by negative FC target stimulation relative to their control targets (**Figure 4B**, left) as well as the pattern resulting from the comparison of negative versus positive FC target stimulation (both relative to their controls; **Figure 4C**) significantly resembled the convergent depression circuit (*r* = 0.50, *p*_Bonf_ < .001 and *r* = 0.51, *p*_Bonf_ < .001, respectively) (47). In contrast, stimulating positive FC targets relative to their controls resulted in a pattern anticorrelated to the depression circuit (*r* = −0.28, *p*_Bonf_ = .005) (**Figure 4D**). None of the condition-specific maps alone was significantly related to the circuit after correction (all *p*_Bonf_ ≥ .19).

## 4 Discussion

In the present study, N = 20 healthy participants received single-pulse TMS at two individually defined left PFC targets exhibiting strong negative and positive FC with sgACC/vmPFC, respectively, as well as two nearby control targets with weaker sgACC/vmPFC-FC but matching peripheral co-stimulation experience, while concurrently measuring TMS-related BOLD changes across the brain. Coil placements were individually optimized to maximize on-target and minimize off-target stimulation based on E-field simulations. This experimental design aimed to assess (i) whether precisely stimulating an individually defined FC target produces downstream sgACC/vmPFC modulation that is specific to that target, beyond what results from stimulating its immediate cortical surroundings (target specificity), and (ii) whether any such specificity depends on the network connectivity profile (positive vs. negative sgACC/vmPFC-FC) of the targeted node (network specificity).

Consistent with prior work combining TMS with concurrent fMRI (e.g., 24,23,26,31), we observed significant TMS-related remote activations within large-scale brain networks, well beyond the immediate prefrontal stimulation sites. Across conditions, positive BOLD modulations were consistently found in the auditory and cingulo-opercular/action mode networks (CON/AMN), as well as in subcortical regions (e.g., thalamus). Negative BOLD modulations were consistently observed in the somatomotor network.

The fact that many of these patterns were consistent across all four stimulation conditions, including control conditions, indicates that a substantial proportion of TMS-related brain-wide effects is attributable to non-specific peripheral co-stimulation (i.e., audible click sounds and scalp sensations) rather than the specific cortical target being engaged (51). The observed pattern closely resembles domain-general responses to salient events within an insula-cingulate network (52), underscoring the necessity of matched control conditions for isolating target-specific downstream TMS effects (53–55).

Critically, stimulation of individually defined DLPFC clusters with strong negative FC to sgACC/vmPFC produced significantly stronger negative BOLD modulations in sgACC, vmPFC, and dorsomedial PFC than stimulation of their nearby control targets with less pronounced sgACC/vmPFC-FC. Importantly, this difference cannot be explained by somatosensory and auditory co-stimulation confounds, which were matched across conditions and additionally included as covariates of no interest in the group-level analyses. Downstream sgACC/vmPFC modulation may therefore be specifically attributable to engaging the individually defined FC target rather than merely a consequence of stimulating the broader cortical region. Furthermore, unlike the condition-specific BOLD patterns alone, the whole-brain pattern of this target-specific effect significantly resembled a convergent depression circuit (47), additionally supporting the potential clinical relevance of specific and selective engagement of negative FC targets.

Relatedly, more negative BOLD responses for precise FC target stimulation may reflect stronger net suppression of downstream neuronal activity in sgACC/vmPFC, thus offering a potential mechanistic explanation for the influential observation that stimulating depressed patients closer to their individually optimal left DLPFC spot with negative sgACC-FC tends to result in better clinical outcomes (10–12). While negative BOLD responses have indeed been related to a decrease in local neuronal activity (e.g., 56), a direct association between BOLD decreases and reduced neuronal activity remains debated and cannot be directly attributed to local inhibitory processes (57,58). Nevertheless, this interpretation is in line with recent studies combining DLPFC-TMS with intracranial EEG recordings and even single-unit recordings in humans and reporting a suppression of high-frequency activity in sgACC (59,60) and excitatory neurons in limbic regions (61), respectively.

In contrast, stimulation of individually defined dorsomedial PFC clusters with strong positive FC to sgACC/vmPFC did not produce significantly different BOLD modulation in sgACC/vmPFC relative to their matched control targets. Of note, positive FC targets experienced weaker induced E-fields than negative FC targets, owing to their greater distance from the TMS coil (62), potentially producing insufficient impact of the stimulation. Furthermore, positive FC targets and their respective control targets differed less in terms of sgACC/vmPFC-FC strength than negative FC targets and their respective control targets (**Figure S6**), potentially reducing experimental sensitivity for detecting subtle differences in downstream modulation.

Beyond the question of target specificity, we found that negative and positive FC target conditions produced qualitatively different downstream modulation patterns, providing evidence for the network-dependence of TMS effects. Stimulation of negative FC targets (relative to their controls), mainly situated within the cingulo-opercular/action-mode network (CON/AMN; **Figure S4**), produced stronger negative BOLD modulations in dorso- and ventromedial as well as subgenual areas, falling within the CON/AMN-anticorrelated default mode network (DMN). Conversely, stimulation of positive FC targets (relative to their controls), situated within the DMN (**Figure S4**), produced relatively stronger negative BOLD modulations in bilateral parietal areas, falling within the DMN-anticorrelated dorsal attention network (DAN). Furthermore, stimulating negative FC targets (both relative to their controls and relative to positive FC targets and their controls) induced a whole-brain BOLD pattern that was significantly correlated with a convergent depression circuit (47), while stimulating oppositely connected positive FC targets (relative to their controls) induced an opposite pattern anticorrelated to the depression circuit. These findings are therefore consistent with (clinically relevant) antagonistic network effects of TMS (63–68).

At the same time, our results stand in contrast to other TMS-fMRI studies reporting (i) positive, rather than negative, sgACC-BOLD responses for DLPFC-TMS (22,69) or (ii) sites with stronger positive, rather than negative, sgACC-FC to produce stronger negative sgACC-BOLD responses in depressed patients (28,29). These discrepancies may reflect methodological variations or genuine differences between healthy participants and depressed patients in sgACC connectivity architecture. This furthermore adds to the broader debate about whether, under what conditions, and for which (deep brain) targets negative or positive FC constitutes the more promising basis for individualized TMS network targeting (9) and highlights that the relationship between a target site’s resting-state functional connectivity profile and its evoked downstream (BOLD) response may not be straightforward.

Some limitations warrant consideration. First, sample size was relatively small (N = 20) and results should be interpreted accordingly. However, this is common for concurrent TMS-fMRI studies (26,31,70), and the well-controlled within-subjects design (totalling almost 80 TMS-fMRI runs) substantially increases statistical power and interpretability. Second, our sample comprised only healthy young participants, and it remains to be established whether findings generalize to clinical populations such as depressed patients, in which FC architecture and baseline sgACC activity may differ (28). This also relates to the state-dependency of TMS effects across the brain (e.g., 71–77) and in PFC (78) and sgACC in particular (79), which we have not explicitly addressed in the present study. Third, realized E-field strengths within FC target clusters were mostly below the target intensity of 100 V/m (38–40,6). Still, detectable effects have been reported for lower E-field strengths within the range achieved in the present study (50-80 V/m) (80,81). Lastly, we did not observe clear BOLD effects at the site of direct cortical stimulation, even though our surface MR coils were specifically designed to maximize signal-to-noise ratio directly below the TMS coil (41,82). However, this is a common observation in concurrent TMS-fMRI studies and may be related to various factors discussed in detail in (70).

In summary, using individually optimized, FC- and E-field-guided precision TMS combined with concurrent fMRI, we provide causal evidence that precisely and selectively stimulating an individually defined cortical FC target produces downstream modulation that may be specifically attributable to the connectivity profile of the targeted site and consistent with anticorrelated network architectures. An important next step will be to test whether these specificity effects are present in clinical populations and whether the magnitude of FC target-specific downstream modulation predicts treatment response to rTMS (cf. 29), thereby closing the mechanistic loop between the network-specific effects of TMS demonstrated here and the clinical outcomes that motivate FC-guided targeting in the first place. More broadly, these findings provide mechanistic support for the premise underlying personalized, FC-guided TMS targeting and precision circuit neuromodulation in general.

## Author contributions

Maximilian Lueckel: Conceptualization, Data curation, Formal analysis, Investigation, Methodology, Project administration, Software, Validation, Visualization, Writing – original draft, Writing – review and editing; Angela Radetz: Investigation, Methodology, Writing – review and editing; Umair Hassan: Methodology, Writing – review and editing; Kenneth Yuen: Methodology, Writing – review and editing; Florian Müller-Dahlhaus: Conceptualization, Writing – review and editing; Raffael Kalisch: Conceptualization, Funding acquisition, Resources, Supervision, Writing – review and editing; Til Ole Bergmann: Conceptualization, Funding acquisition, Project administration, Resources, Supervision, Writing – review and editing.

