## Supplement for "Target- and network-specific downstream effects of personalized precision transcranial magnetic stimulation: Evidence from concurrent functional neuroimaging"

Maximilian Lueckel[1,2,†], Angela Radetz[2], Umair Hassan[1,2,3,4], Kenneth Yuen[1,2], Florian Müller-Dahlhaus[5,2], Raffael Kalisch[2,1,*], Til Ole Bergmann[2,1,*]

[1] Leibniz Institute for Resilience Research (LIR) gGmbH, Mainz, Germany

[2] Neuroimaging Center (NIC), Focus Program Translational Neuroscience (FTN), Johannes Gutenberg University Medical Center, Mainz, Germany

[3] Department of Psychiatry and Behavioral Sciences, Stanford University, Palo Alto, CA, USA

[4] Stanford Center for Sleep and Circadian Sciences, Stanford University, Palo Alto, CA, USA

[5] Department of Psychiatry and Psychotherapy, Johannes Gutenberg University Medical Center, Mainz, Germany

*contributed equally

### Supplementary Methods

#### Experimental procedure

*Intake session.* An intake MRI session comprised resting-state functional and anatomical scans (see *MRI data acquisition and processing* below). Head motion was restricted using foam pads and tape attached to participants' forehead and MR coil housing. During resting-state scans, participants were instructed to keep their eyes open and stay awake, fixate a light grey cross on a black background (viewed via a mirror on the MR coil), and let their mind wander (i.e., not to think of anything particular or stick to one specific thought). Alertness was monitored via eye-tracking, and participants rated their tiredness after each run (0 = not tired at all/wide awake, 10 = extremely tired/almost falling asleep). Participants were allowed to close their eyes during the anatomical scans.

*Concurrent TMS-fMRI sessions.* Each participant underwent two TMS-fMRI sessions, each comprising two runs (one participant completed only one session (two runs) and another participant completed only three runs due to technical difficulties). Each session targeted either the left *dorsolateral* PFC target with strong *negative* FC to sgACC/vmPFC (and its control) or the left *dorsomedial* PFC target with strong *positive* FC to sgACC/vmPFC (and its control; see below for target selection). FC and control target order was counterbalanced within sessions, and positive/negative FC target order was counterbalanced across sessions, which were separated by at least 2 days.

At the start of each session, the participant was positioned on the MR table, and optical neuronavigation trackers were attached to their forehead and to an air-cooled MR-compatible figure-of-eight TMS coil (MRi-B91, MagVenture, Denmark) (**Figure 1G** in the main text). An MR-compatible stereo infrared camera recorded these trackers, which, combined with the Localite neuronavigation system (TMS-Navigator-MR, Localite, Germany), enabled precise MR-based navigation of the TMS coil to individually optimized placements (see *TMS targeting optimization* below) and, whenever possible, continuous monitoring of coil placement inside the scanner bore during fMRI.

After confirming that both FC and control coil placements were reachable, we set the stimulation intensity. Titration always began with the FC target placement: the TMS coil was held in place by one experimenter while another gradually increased the intensity from 30% MSO in 5% steps, applying several single pulses (3-5 s inter-pulse interval) at each step while the participant continuously indicated tolerability. Where possible, intensity was raised to the individually optimized value derived from E-field simulations (see below). Otherwise, we used the maximum tolerable intensity. At the selected intensity, 4-5 further pulses were applied, and the participant rated stimulation unpleasantness (0 = not unpleasant at all, 10 = extremely unpleasant/almost unbearable) and memorized the subjective experience of the stimulation for comparison.

We then repeated this titration procedure for the control coil placement (again starting at 30% MSO), but here the participant indicated the intensity at which the subjective experience (unpleasantness, loudness, peripheral co-stimulation) matched that of the FC target stimulation as closely as possible. This intensity was used for the control target. This procedure both calibrated stimulation intensity to each participant's tolerability and familiarized them with TMS in the scanner environment while in a supine position.

The participant's head was then stabilized with a vacuum cushion, and the TMS coil was navigated to the first run's placement (FC or control target) and fixed with a dedicated MR-compatible coil holder (**Figure 1G** in the main text; MagVenture, Denmark) to prevent movement during fMRI. Participants were equipped with an emergency bell and stop switch for the TMS device before being moved into the scanner isocenter.

In each run, we applied 65 single biphasic TMS pulses (to allow reliable assessment of TMS-evoked BOLD responses (1)) at the individually adjusted intensity (see *Determination of stimulation intensity* below), with a jittered inter-pulse interval of 10-15 s. Before each pulse at target intensity, two low-intensity, sub-threshold pulses (~35% of target intensity) (**Figure S1; Table S1**) were applied to prepare participants for the stronger pulse and thereby reduce startle-related head movements and eye blinks (with an inter-pulse interval corresponding to the TR of the respective EPI sequence).

The TMS device (MagPro X100, MagVenture, Denmark) was positioned outside the MR room and connected to the coil via an extension cable and high-voltage low-pass filter, controlled by the BEST toolbox (2), which synchronized TMS application to MR volume triggers. Pulses were applied in the gap between MR volume acquisitions, starting 10 ms after and ending at least 144 ms before adjacent acquisitions to avoid imaging artifacts from electromagnetic interference and coil vibration (3).

Immediately after each run, participants rated via intercom (i) stimulation unpleasantness (0-10), (ii) loudness of the TMS click (0-10), (iii) presence, location, and strength of any TMS-related muscle twitches (0-10), and (iv) tiredness during the measurement (0-10).

#### TMS targeting optimization

_Functional connectivity analysis._ Individual FC maps for target identification and coil placement optimization were computed from participants' preprocessed intake MRI data (see *MRI Data Acquisition and Processing* below) using a surface-based seed map method (4,5). A normative, whole-brain sgACC-/vmPFC-based FC map was derived from the dense connectome of a large HCP sample (N = 1003) (6), using the mean time series from two 10 mm-radius ROIs centered on previously reported sgACC (X = 5.9, Y = 16.3, Z = -9.8) (7) and vmPFC (X = 6, Y = 46, Z = -14) (8) coordinates (**Figure S2A**). This normative map (excluding the left dorsal PFC, which encompassed our TMS targets; **Figure 1A** in the main text**, Figure S2B-C**; following (4,5)) was then used as a weighted seed in a whole-brain FC analysis of each participant's resting-state data (**Figure 1B** in the main text). This seed map approach yields more reliable single-subject connectivity estimates than using a small ROI as seed (5,9,4).

_TMS target identification._ We first multiplied the individual FC maps vertex-wise with individual FreeSurfer-derived sulcal depth maps, upweighting FC clusters on the gyral crown (**Figure S3**; cf. (10)), where the TMS-induced E-field is maximal (11,12). Following the cluster-based approach of (5), these sulcal depth-weighted FC maps were restricted to the left dorsal PFC (**Figure 1C** in the main text), within which we identified the 5% of vertices with the strongest negative and positive sgACC/vmPFC-FC. The largest resulting negative and positive clusters were selected as target clusters of interest (**Figure 1C** in the main text) and iteratively expanded or constricted to ~50 surface vertices, to avoid biases in cluster size (and thus required stimulation intensity) across participants and stimulation conditions (see *Determination of stimulation intensity* below). Combining seed map and cluster-based methods yields more reliable individual-level FC-based TMS targets than using a small ROI as seed and/or the peak (anti)correlation vertex/voxel (4,5).

_Electric field modelling._ We used SimNIBS (version 4.0) to simulate TMS-induced E-fields (13) and individually determine the optimal coil placement for stimulating each participant's negative and positive FC target clusters. A realistic volume conductor model of each participant's head was created by segmenting their anatomical MR image into tissue types (grey and white matter, CSF, bone, skin, etc.) with standard conductivity values, using the *charm* pipeline (14). E-field simulations were run for up to 770 potential coil placements, comprising a search grid of up to 70 positions (25 mm radius, 5 mm steps) and 11 orientation angles per position (180°, 15° steps) (15), centered on the point on the head surface closest to the center of gravity of each FC cluster (**Figure 1D** in the main text).

After identifying the optimal coil position and orientation for each FC cluster (selection criteria below), we ran a second set of E-field simulations to determine the corresponding control placements. Here, we generated a search grid (same angle resolution) surrounding the optimized position (40 mm radius, 5 mm steps), excluding positions within a 10 mm radius of the optimized position and the previous grid's center (**Figure 1D** in the main text), to prevent control placements from being too close to the FC-optimized placement and effectively co-stimulating the FC target cluster.

_Selection of optimal TMS coil placements._ The optimal coil placement for each FC cluster was the combination of position and orientation that maximized both E-field magnitude within the target cluster and overlap between the whole-brain sgACC/vmPFC FC map and the E-field map (by minimizing or maximizing the sum of vertex-wise FC-by-E-field products, respectively) (**Figure 1E** in the main text). Unlike existing targeting procedures (e.g., SimNIBS's auxiliary dipole method), which treat the cortical target as a single coordinate or as a spatially restricted ROI within which E-field strength is maximized, this approach additionally accounts for the whole-brain topography of both FC and induced E-field. This ensures the E-field maximally affects target areas (expressing negative or positive sgACC/vmPFC-FC) while largely sparing off-target areas (e.g., of opposite FC valence; **Figure 1E** in the main text).

Control placements were determined similarly. We first "inverted" each participant's FC map, so that areas with strong negative or positive sgACC/vmPFC-FC showed minimal values and near-zero-FC areas showed maximal values. As there was no dedicated target cluster here, we simply maximized overlap (i.e., the sum of vertex-wise products) between the inverted FC map and the E-field map, sparing areas of negative and positive sgACC/vmPFC-FC as much as possible. We then selected the coil position/orientation that would (i) stimulate cortex with a mixed (i.e., less specific) and thus on average weaker sgACC/vmPFC-FC profile than the corresponding target cluster, preserving connectivity specificity to the FC target; (ii) lie as close as possible to the FC-optimized placement, to match peripheral (somatosensory and auditory) co-stimulation, a serious confound in any TMS study (16,17); and (iii) induce an E-field as weak as possible within the corresponding FC target cluster, to avoid effectively co-stimulating it. This approach aimed to simultaneously maximize experimental variance and minimize confounding variance between conditions.

*Determination of stimulation intensity*. Stimulation intensity was individually determined from the same E-field simulations, aiming for a mean E-field strength of 100 V/m within the FC target clusters (18–20,10). The corresponding di/dt value (rate of change of coil current) was translated into % maximum stimulator output (MSO) value for our setup. This simulation-derived intensity could not always be realized, for two reasons: (i) technical limits: the required MSO exceeded 100% (e.g., when the target was too deep, exacerbated by power loss from the long cable and added coil-to-head distance from the MR coil) (3); or (ii) tolerability: the participant could not tolerate the derived intensity. In both cases, we used the highest technically possible (100% MSO) or tolerable intensity (see *Experimental Procedure* for details).

### MRI data acquisition and processing

#### Intake MRI data

*Acquisition.* During the intake session, using a 32-channel Siemens head coil in a Siemens Magnetom Prisma 3T scanner at the Neuroimaging Center (NIC) of the University Medical Center Mainz, we acquired ~15 minutes (610 volumes) of resting-state fMRI using a T2*-weighted multi-band, multi-echo (MBME) EPI sequence (repetition time (TR): 1435 ms; echo time $(TE)_{1/2/3/4}$: 12/28.24/44.48/60.72 ms; field-of-view (FOV): 215 mm; flip angle (FA): 65°; 2.5 mm isotropic voxels; 51 slices; anterior-posterior (AP) phase encoding direction; in-plane acceleration factor: 2; multi-band (MB) acceleration factor: 3). Notably, multi-echo fMRI has been shown to yield more reliable single-subject FC estimates than single-echo data (21–24), which is essential for FC-based TMS target selection (5,4). We then acquired high-resolution MPRAGE T1-weighted (TR: 2500 ms; TE: 2.22 ms; FOV: 256 mm; FA: 8°) and T2-weighted (TR: 3200 ms; TE: 563 ms; FOV: 256 mm) anatomical images (0.8 mm isotropic voxels; 208 sagittal slices) for MR-based neuronavigation and head modelling for E-field simulations.

Preprocessing. Resting-state fMRI preprocessing closely followed (21). Briefly, T1- and T2-weighted images were processed with the Human Connectome Project anatomical pipeline (25), including ACPC-alignment, brain extraction, and MNI registration using FSL (26) and FreeSurfer (27). FreeSurfer additionally generated cortical surfaces for later surface-based analyses. Functional processing included co-registration of each echo to the ACPC-aligned T1w image and correction for spatial distortions, slice timing, and motion (FSL). Echoes were then optimally combined and submitted to multi-echo independent component analysis (ME-ICA) using the *tedana* workflow (version 0.0.11) (28), which classifies components as BOLD-like signal (retained) or non-BOLD noise (discarded) based on their decay across echoes (29,30). Components were also visually reviewed and reclassified if necessary, following (31) and (32). Optimally combined images were denoised by regressing out noise-component time series and global grey matter signal and subsequently bandpass-filtered (0.01-0.1 Hz). After MNI transformation, data were mapped to the individual cortical surface and combined into CIFTI format, representing surface and volume data simultaneously (25), and finally spatially smoothed (geodesic σ = 2.58 mm; Euclidean FWHM = 6 mm). See (21) for details.

**TMS-fMRI data**

Acquisition. For concurrent TMS-fMRI, we used two TMS-compatible ultrathin 7-channel RF surface coil arrays (MagVenture, Denmark) (33,34) for image acquisition: one attached to the TMS coil (between the head and coil), the other on the right lateral head surface (for whole-brain coverage; **Figure 1G** in the main text). The T2*-weighted MBME EPI sequence from the intake session (see *Intake MRI data* above) was adapted for these coils and interleaved TMS application. Sequence parameters were slightly adjusted during the study to improve image quality. For n = 6 participants, we used the following parameters: 1095 ms TA + 155 ms gap between volumes for interleaved TMS = 1250 ms TR; $TE_{1/2/3}$: 13/34.62/56.24 ms; FOV: 215 mm; FA: 48°; 3.2 mm isotropic voxels; 39 slices; AP phase encoding direction; MB acceleration factor: 3; 7/8 partial Fourier. For n = 14 participants, the following parameters were changed: 1137 ms TA + 163 ms gap between volumes for interleaved TMS = 1300 ms TR; $TE_{1/2/3}$: 14.6/36.22/57.84 ms; FA: 65°.

Preprocessing. Preprocessing the concurrent TMS-fMRI data required additional steps, mainly due to pronounced image inhomogeneity from the MR surface coil design (35). NORDIC denoising was applied to each run and echo individually to reduce thermal noise (using magnitude images and a theoretical noise level) (36,37). We then applied six iterations of ANTs bias field correction (38–42). Since skin/fat signal directly below the coils appeared brighter and posed a challenge for, e.g., co-registration algorithms, we masked out non-brain structures using (dilated) brain masks derived using SynthStrip (43) before further processing.

fMRIPrep (version 25.1.1) (44) was then used to preprocess the individual echoes. An EPI reference volume was generated from the shortest echo of each run using fMRIPrep's custom methodology. Fieldmaps were estimated per run from two EPI references with opposing phase-encoding directions using *topup* (45) and rigidly registered to the EPI reference. Motion correction used *mcflirt* (FSL), realigning all volumes to the EPI reference, and slice timing correction used *3dTShift* (AFNI) (46). The EPI reference was co-registered to the T1w reference using *mri_coreg* (FreeSurfer), followed by FSL's *flirt* (47) with boundary-based registration (48) and six degrees of freedom. Functional time series were then projected to the individual cortical surface and resampled to symmetric "fsLR" space using Connectome Workbench (25), excluding voxel time series with locally high coefficient of variation.

Like the resting-state data above, fMRIPrep-preprocessed functional echoes (volume space) were submitted to the *tedana* workflow (version 25.0.1) for ME-ICA and optimal echo combination (28). Components were automatically classified as BOLD (TE-dependent) or non-BOLD (TE-independent) using the tedana_orig decision tree (similar to the MEICA v2.5 tree of (30); https://tedana.readthedocs.io/en/stable/included_decision_trees.html), but also visually reviewed and reclassified as needed, following (31), (32), and (49), with particular attention to TMS-related motion or electromagnetic interference artifacts that could bias the subsequent GLM analysis (49). Rejected components' time series were orthogonalized to the accepted components and, next to unstructured noise time series from the whole brain, white matter, and CSF, regressed from the data.

The optimally combined and denoised data were spatially normalized to MNI space via nonlinear ANTs registration (for group-level volume-space analyses) and mapped to the individual cortical surface. Whole-brain volume and surface data were then smoothed with Euclidean (FWHM = 6 mm) and geodesic ($\sigma$ = 2.58 mm) Gaussian kernels, respectively.

*Statistical analysis.* Nilearn (version 0.10.4) (50) in Python (3.11) and SPM25 in Matlab (R2022a) were used for statistical analysis of the preprocessed TMS-fMRI data. At the first (single-subject) level, each TMS pulse was modelled as a single instantaneous event, parametrically modulated by the realized % MSO and convolved with SPM's canonical HRF. Next to the noise time series mentioned above, the design matrix also included the six motion parameters with their temporal derivatives and quadratic expansions as nuisance covariates and five polynomial regressors for high-pass filtering. This design matrix was used in a voxel-wise GLM with generalized least squares and a global approximate AR(1) model. We then computed the contrast for the TMS pulse regressor, yielding one first-level contrast map per stimulation condition (negative FC target, negative FC *control* target, positive FC target, positive FC *control* target) per participant.

**Inclusion and exclusion criteria**

Inclusion criteria:

- Age between 18 and 45 years.
- Good physical and mental health.
- Understanding of study procedures and agreement to participate in the study by giving written informed consent.
- Willingness to comply with the study restrictions.

Exclusion criteria:

- Younger than the age of legal consent.
- Mental or legal incapacitance, significant emotional problems or history of a clinically significant psychiatric disorder (as defined by ICD-10).
- History of any illness that, in the opinion of the study investigator, might confound the results of the study or pose an additional risk to the participant by their participation in the study.
- History of stroke, seizures, or major neurological disorder.
- History of significant head injury/trauma with loss of consciousness lasting for ≥15 minutes and one or more of the following:
    - Recurring seizures resulting from the head injury
    - Persistent neurological or cognitive sequelae of the injury
    - Cognitive rehabilitation following the injury
- Family history of epilepsy.
- Cardiac pacemaker, implanted medication pump, intracardiac line, or acute, unstable cardiac disease.
- Intracranial implant (e.g., aneurysm clips, shunts, stimulators, cochlear implants, or electrodes) or any other metal object within or near the head (excluding the mouth) that cannot be safely removed.
- Regular use (including “recreational use”) of any illicit drugs or a history of drug (including alcohol) abuse.
- History of low blood pressure and/or a history of repeated hypotensive faints.
- Pregnancy or breastfeeding or trying to get pregnant.
- Any contraindication to MRI or TMS.

### Supplementary Results

#### TMS targets and coil placements

Relative to the average coil placement (black coil representations in **Figure 2B**), the mean Euclidean distance of individual coil positions was M = 10.11 mm (SD = 5.90 mm) for negative and M = 23.44 mm (SD = 16.09 mm) for positive FC targets ($t(18) = -3.35$, $p = .004$, $d = -1.09$; the larger deviation for positive targets likely reflecting the greater surface extent of the DMN across the prefrontal cortex). Absolute deviations of individually optimized coil orientations from the average were M = 25.8° (SD = 21.9°) for negative and M = 35.5° (SD = 35.6°) for positive FC targets (**Figure S5**).

Coil position centers for FC targets and their respective control targets were on average 30.33 mm apart (SD = 6.97 mm) for negative and 31.91 mm apart (SD = 5.52 mm) for positive conditions, which did not differ significantly between conditions (Wilcoxon $V = 72$, $p = .580$, $r = -0.16$ [−0.60, 0.35], n = 18).

#### Confounds

Negative FC targets and controls, positive FC targets and controls, and negative and positive FC targets were not significantly different from each other (all $p_{\text{Bonf}} \geq .078$) regarding stimulation intensity (% MSO; negative FC target vs. control: mean difference = 2.10 [−3.56, 7.76], Wilcoxon $V = 76.5$, $p_{\text{Bonf}} > .99$, $r = 0.28$ [−0.40, 0.76]; negative vs. positive FC target: mean difference = −4.84 [−11.69, 2.01], $t(18) = -1.87$, $p_{\text{Bonf}} = .235$, $d = -0.35$ [−0.83, 0.13]; positive FC target vs. control: mean difference = −0.56 % [−7.21, 6.10], Wilcoxon $V = 25.5$, $p_{\text{Bonf}} > .99$, $r = -0.23$ [−0.78, 0.53]), subjectively reported unpleasantness (negative FC target vs. control: mean difference = 0.12 [−0.81, 1.06], $t(19) = 0.35$, $p_{\text{Bonf}} > .99$, $d = 0.06$ [−0.35, 0.47]; negative vs. positive FC target: mean difference = 0.32 [−1.48, 2.11], $t(18) = 0.47$, $p_{\text{Bonf}} > .99$, $d = 0.14$ [−0.64, 0.93]; positive FC target vs. control: mean difference = 0.44 [−1.19, 2.08], Wilcoxon $V = 46.0$, $p_{\text{Bonf}} > .99$, $r = 0.18$ [−0.54, 0.75]) and subjectively reported loudness of the TMS click sound (negative FC target vs. control: mean difference = 0.03 [−0.47, 0.52], $t(19) = 0.13$, $p_{\text{Bonf}} > .99$, $d = 0.02$ [−0.29, 0.32]; negative vs. positive FC target: mean difference = 0.08 [−1.04, 1.19], $t(18) = 0.19$, $p_{\text{Bonf}} > .99$, $d = 0.05$ [−0.64, 0.75]; positive FC target vs. control: mean difference = 0.78 [−0.07, 1.62], $t(17) = 2.44$, $p_{\text{Bonf}} = .078$, $d = 0.53$ [−0.05, 1.11])

**Table S1**. Descriptive statistics (mean and standard deviation) of mean E-field strength in individual FC clusters for low-intensity pre-pulses and mean FC strength in E-field hotspots, for each stimulation condition.

| | **Stimulation condition** | | | | | | | |
|---|---|---|---|---|---|---|---|---|
| | **Negative FC control target (n = 20)** | | **Negative FC target (n = 20)** | | **Positive FC target (n = 19)** | | **Positive FC control target (n = 18)** | |
| | **M** | **SD** | **M** | **SD** | **M** | **SD** | **M** | **SD** |
| **Mean E-field [V/m] values in FC clusters for low-intensity pulses (cf. Figure S1)** | | | | | | | | |
| Negative FC cluster | 10.86 | 4.64 | 28.15 | 6.19 | 7.65 | 4.85 | 9.42 | 2.83 |
| Positive FC cluster | 7.63 | 2.59 | 9.65 | 3.95 | 24.93 | 6.75 | 12.70 | 2.62 |
| **Mean FC values in E-field hotspots (cf. Figure S6)** | | | | | | | | |
| | 0.11 | 0.13 | -0.20 | 0.08 | 0.48 | 0.09 | 0.40 | 0.16 |

*Note.* Means and SDs are based on all available data per condition, while statistical tests only used complete pairs.

**Table S2**. Second-level GLM results for the [Negative FC control target > Baseline] contrast.

| cluster-level | | | | peak-level | | | | | coordinates | | |
|---|---|---|---|---|---|---|---|---|---|---|---|
| $p_{FWE-corr}$ | $q_{FDR-corr}$ | $k_E$ | $p_{uncorr}$ | $p_{FWE-corr}$ | $q_{FDR-corr}$ | T | $(Z_E)$ | $p_{uncorr}$ | X | Y | Z |
| **Negative FC *control* target > Baseline** | | | | | | | | | | | |
| 0.000 | 0.000 | 8140 | 0.000 | 0.000 | 0.003 | 11.68 | 6.04 | 0.000 | 66 | -22 | 18 |
| | | | | 0.001 | 0.003 | 10.85 | 5.86 | 0.000 | 54 | -30 | 18 |
| | | | | 0.001 | 0.004 | 10.15 | 5.69 | 0.000 | 56 | 6 | 2 |
| 0.000 | 0.000 | 5250 | 0.000 | 0.000 | 0.003 | 11.17 | 5.93 | 0.000 | -36 | -16 | -2 |
| | | | | 0.006 | 0.010 | 9.21 | 5.45 | 0.000 | -38 | -6 | -6 |
| | | | | 0.008 | 0.011 | 9.01 | 5.39 | 0.000 | -40 | -24 | -2 |
| 0.000 | 0.000 | 233 | 0.000 | 0.211 | 0.036 | 6.98 | 4.73 | 0.000 | 6 | -56 | -14 |
| | | | | 0.723 | 0.114 | 5.82 | 4.26 | 0.000 | -12 | -66 | -16 |
| | | | | 0.996 | 0.292 | 4.85 | 3.79 | 0.000 | -4 | -68 | -12 |
| 0.000 | 0.000 | 654 | 0.000 | 0.480 | 0.072 | 6.27 | 4.45 | 0.000 | 4 | 4 | 64 |
| | | | | 0.786 | 0.127 | 5.69 | 4.20 | 0.000 | -10 | 6 | 34 |
| | | | | 0.983 | 0.246 | 5.05 | 3.89 | 0.000 | 0 | 22 | 36 |
| 0.001 | 0.000 | 222 | 0.000 | 0.767 | 0.126 | 5.73 | 4.22 | 0.000 | 6 | -36 | -46 |
| | | | | 0.933 | 0.184 | 5.32 | 4.03 | 0.000 | -6 | -36 | -30 |
| | | | | 0.962 | 0.205 | 5.20 | 3.97 | 0.000 | -4 | -36 | -46 |

**Table S3**. Second-level GLM results for the [Negative FC control target < Baseline] contrast.

| cluster-level | | | | peak-level | | | | | coordinates | | |
|---|---|---|---|---|---|---|---|---|---|---|---|
| $p_{FWE\text{-}corr}$ | $q_{FDR\text{-}corr}$ | $k_E$ | $p_{uncorr}$ | $p_{FWE\text{-}corr}$ | $q_{FDR\text{-}corr}$ | T | $(Z_E)$ | $p_{uncorr}$ | X | Y | Z |
| **Negative FC *control* target < Baseline** | | | | | | | | | | | |
| 0.000 | 0.000 | 3227 | 0.000 | 0.011 | 0.131 | 8.80 | 5.33 | 0.000 | 18 | -62 | 68 |
| | | | | 0.07 | 0.157 | 7.72 | 4.99 | 0.000 | 24 | -64 | 62 |
| | | | | 0.178 | 0.181 | 7.11 | 4.78 | 0.000 | -18 | -70 | 62 |
| 0.000 | 0.000 | 1021 | 0.000 | 0.116 | 0.157 | 7.43 | 4.9 | 0.000 | 46 | -22 | 42 |
| | | | | 0.401 | 0.213 | 6.44 | 4.52 | 0.000 | 36 | -30 | 62 |
| | | | | 0.761 | 0.245 | 5.74 | 4.23 | 0.000 | 34 | -12 | 64 |
| 0.000 | 0.000 | 333 | 0.000 | 0.340 | 0.213 | 6.58 | 4.58 | 0.000 | -30 | -84 | 26 |
| | | | | 0.999 | 0.458 | 4.71 | 3.72 | 0.000 | -46 | -78 | 30 |
| | | | | 1.000 | 0.502 | 4.55 | 3.63 | 0.000 | -38 | -88 | 22 |
| 0.000 | 0.000 | 1276 | 0.000 | 0.414 | 0.213 | 6.41 | 4.51 | 0.000 | -42 | -38 | 54 |
| | | | | 0.523 | 0.216 | 6.19 | 4.42 | 0.000 | -38 | -42 | 66 |
| | | | | 0.614 | 0.233 | 6.02 | 4.35 | 0.000 | -50 | -28 | 46 |
| 0.000 | 0.000 | 289 | 0.000 | 0.469 | 0.213 | 6.29 | 4.46 | 0.000 | -44 | -80 | -10 |
| | | | | 0.679 | 0.241 | 5.90 | 4.29 | 0.000 | -42 | -68 | -6 |
| | | | | 1.000 | 0.821 | 3.91 | 3.25 | 0.001 | -38 | -68 | -14 |
| 0.001 | 0.000 | 205 | 0.000 | 0.488 | 0.213 | 6.26 | 4.45 | 0.000 | 54 | -68 | -6 |
| | | | | 0.999 | 0.463 | 4.66 | 3.69 | 0.000 | 42 | -70 | -8 |
| 0.000 | 0.000 | 474 | 0.000 | 0.499 | 0.213 | 6.24 | 4.44 | 0.000 | 10 | 24 | 14 |
| | | | | 0.675 | 0.241 | 5.91 | 4.30 | 0.000 | 14 | 18 | 18 |
| | | | | 0.856 | 0.249 | 5.54 | 4.13 | 0.000 | 22 | 38 | 12 |
| 0.001 | 0.000 | 218 | 0.000 | 0.595 | 0.233 | 6.05 | 4.36 | 0.000 | 0 | 56 | -2 |
| | | | | 1.000 | 0.678 | 4.15 | 3.40 | 0.000 | 4 | 64 | 8 |
| | | | | 1.000 | 0.935 | 3.73 | 3.15 | 0.001 | 10 | 56 | 8 |
| 0.043 | 0.008 | 102 | 0.001 | 0.770 | 0.245 | 5.73 | 4.22 | 0.000 | 24 | -50 | -14 |
| | | | | 0.998 | 0.437 | 4.78 | 3.75 | 0.000 | 24 | -64 | -8 |
| 0.000 | 0.000 | 233 | 0.000 | 0.821 | 0.248 | 5.62 | 4.17 | 0.000 | 28 | 6 | 36 |
| | | | | 0.837 | 0.248 | 5.59 | 4.15 | 0.000 | 24 | -2 | 28 |
| 0.002 | 0.000 | 179 | 0.000 | 0.885 | 0.268 | 5.47 | 4.10 | 0.000 | -22 | 10 | 30 |
| | | | | 0.979 | 0.351 | 5.08 | 3.91 | 0.000 | -20 | 18 | 36 |
| | | | | 0.999 | 0.463 | 4.68 | 3.70 | 0.000 | -20 | 30 | 18 |
| 0.002 | 0.000 | 182 | 0.000 | 0.987 | 0.369 | 5.00 | 3.87 | 0.000 | 0 | 58 | 24 |
| | | | | 1.000 | 0.603 | 4.34 | 3.51 | 0.000 | 6 | 56 | 44 |
| | | | | 1.000 | 0.603 | 4.32 | 3.50 | 0.000 | -6 | 58 | 38 |

**Table S4**. Second-level GLM results for the [Negative FC target > Baseline] contrast.

| cluster-level | | | | peak-level | | | | | coordinates | | |
|---|---|---|---|---|---|---|---|---|---|---|---|
| $p_{FWE\text{-}corr}$ | $q_{FDR\text{-}corr}$ | $k_E$ | $p_{uncorr}$ | $p_{FWE\text{-}corr}$ | $q_{FDR\text{-}corr}$ | T | $(Z_E)$ | $p_{uncorr}$ | X | Y | Z |
| **Negative FC target > Baseline** | | | | | | | | | | | |
| 0.000 | 0.000 | 9345 | 0.000 | 0.000 | 0.005 | 10.98 | 5.89 | 0.000 | 42 | -4 | -4 |
| | | | | 0.003 | 0.013 | 9.67 | 5.57 | 0.000 | 50 | -26 | 22 |
| | | | | 0.005 | 0.014 | 9.34 | 5.48 | 0.000 | 58 | -24 | 22 |
| 0.000 | 0.000 | 5666 | 0.000 | 0.017 | 0.014 | 8.57 | 5.26 | 0.000 | -42 | -4 | -6 |
| | | | | 0.057 | 0.024 | 7.83 | 5.03 | 0.000 | -54 | -14 | 18 |
| | | | | 0.173 | 0.037 | 7.09 | 4.77 | 0.000 | -40 | -18 | 2 |
| 0.003 | 0.001 | 183 | 0.000 | 0.145 | 0.037 | 7.23 | 4.82 | 0.000 | 52 | 0 | 54 |
| | | | | 1.000 | 0.380 | 4.54 | 3.63 | 0.000 | 46 | -6 | 48 |
| | | | | 1.000 | 0.526 | 4.25 | 3.46 | 0.000 | 38 | -8 | 52 |
| 0.000 | 0.000 | 773 | 0.000 | 0.317 | 0.047 | 6.61 | 4.59 | 0.000 | 6 | -24 | 26 |
| | | | | 0.339 | 0.048 | 6.55 | 4.57 | 0.000 | 12 | -28 | 38 |
| | | | | 0.551 | 0.075 | 6.10 | 4.38 | 0.000 | -6 | -40 | 26 |
| 0.003 | 0.001 | 179 | 0.000 | 0.365 | 0.052 | 6.49 | 4.54 | 0.000 | -32 | -52 | -30 |
| | | | | 0.973 | 0.197 | 5.10 | 3.92 | 0.000 | -28 | -64 | -24 |
| | | | | 0.998 | 0.305 | 4.71 | 3.72 | 0.000 | -26 | -58 | -30 |
| 0.000 | 0.000 | 1278 | 0.000 | 0.626 | 0.086 | 5.96 | 4.32 | 0.000 | 6 | 30 | 36 |
| | | | | 0.826 | 0.128 | 5.58 | 4.15 | 0.000 | 8 | 16 | 34 |
| | | | | 0.841 | 0.131 | 5.54 | 4.13 | 0.000 | -6 | 12 | 34 |
| 0.026 | 0.004 | 119 | 0.001 | 0.693 | 0.098 | 5.84 | 4.27 | 0.000 | -6 | -62 | -8 |
| | | | | 0.936 | 0.162 | 5.27 | 4.01 | 0.000 | 0 | -56 | 0 |
| | | | | 1.000 | 0.411 | 4.47 | 3.59 | 0.000 | 0 | -66 | -14 |
| 0.018 | 0.003 | 129 | 0.001 | 0.913 | 0.152 | 5.35 | 4.04 | 0.000 | 14 | -68 | -20 |
| | | | | 0.943 | 0.166 | 5.25 | 3.99 | 0.000 | 8 | -60 | -8 |
| | | | | 0.999 | 0.312 | 4.69 | 3.71 | 0.000 | 18 | -64 | -26 |

**Table S5**. Second-level GLM results for the [Negative FC target < Baseline] contrast.

| cluster-level | | | | peak-level | | | | | coordinates | | |
|---|---|---|---|---|---|---|---|---|---|---|---|
| $p_{FWE-corr}$ | $q_{FDR-corr}$ | $k_E$ | $p_{uncorr}$ | $p_{FWE-corr}$ | $q_{FDR-corr}$ | T | $(Z_E)$ | $p_{uncorr}$ | X | Y | Z |
| **Negative FC target < Baseline** | | | | | | | | | | | |
| 0.000 | 0.000 | 676 | 0.000 | 0.038 | 0.159 | 8.08 | 5.11 | 0.000 | -62 | -14 | -6 |
| | | | | 0.242 | 0.222 | 6.83 | 4.68 | 0.000 | -62 | 2 | -14 |
| | | | | 0.590 | 0.280 | 6.03 | 4.35 | 0.000 | -52 | 10 | -22 |
| 0.004 | 0.001 | 170 | 0.000 | 0.045 | 0.159 | 7.98 | 5.08 | 0.000 | -48 | -68 | -12 |
| | | | | 1.000 | 0.662 | 4.37 | 3.53 | 0.000 | -50 | -54 | -14 |
| 0.000 | 0.000 | 4979 | 0.000 | 0.066 | 0.159 | 7.76 | 5.01 | 0.000 | 14 | 6 | 24 |
| | | | | 0.075 | 0.159 | 7.68 | 4.98 | 0.000 | 18 | -8 | 28 |
| | | | | 0.220 | 0.222 | 6.91 | 4.70 | 0.000 | 32 | -6 | 26 |
| 0.000 | 0.000 | 863 | 0.000 | 0.111 | 0.178 | 7.44 | 4.90 | 0.000 | -6 | -30 | 76 |
| | | | | 0.619 | 0.280 | 5.97 | 4.33 | 0.000 | 6 | -30 | 70 |
| | | | | 0.627 | 0.280 | 5.96 | 4.32 | 0.000 | -4 | -40 | 76 |
| 0.000 | 0.000 | 293 | 0.000 | 0.344 | 0.246 | 6.54 | 4.56 | 0.000 | 38 | -86 | 2 |
| | | | | 0.997 | 0.537 | 4.77 | 3.75 | 0.000 | 36 | -82 | -12 |
| | | | | 1.000 | 0.599 | 4.57 | 3.64 | 0.000 | 28 | -90 | 10 |
| 0.028 | 0.005 | 117 | 0.001 | 0.486 | 0.280 | 6.22 | 4.43 | 0.000 | -4 | 32 | 62 |
| | | | | 0.996 | 0.527 | 4.81 | 3.77 | 0.000 | -12 | 34 | 54 |
| 0.000 | 0.000 | 240 | 0.000 | 0.502 | 0.280 | 6.19 | 4.42 | 0.000 | 28 | -82 | -32 |
| | | | | 0.969 | 0.407 | 5.12 | 3.93 | 0.000 | 26 | -74 | -28 |
| | | | | 0.996 | 0.526 | 4.83 | 3.78 | 0.000 | 40 | -78 | -40 |
| 0.036 | 0.006 | 111 | 0.001 | 0.571 | 0.280 | 6.06 | 4.37 | 0.000 | -36 | -90 | -0 |
| | | | | 0.757 | 0.280 | 5.72 | 4.21 | 0.000 | -34 | -86 | -10 |
| 0.000 | 0.000 | 342 | 0.000 | 0.591 | 0.280 | 6.02 | 4.35 | 0.000 | -44 | -34 | 68 |
| | | | | 0.734 | 0.280 | 5.76 | 4.23 | 0.000 | -42 | -22 | 60 |
| | | | | 0.978 | 0.416 | 5.06 | 3.90 | 0.000 | -52 | -24 | 52 |
| 0.000 | 0.000 | 453 | 0.000 | 0.720 | 0.280 | 5.79 | 4.25 | 0.000 | -30 | -2 | 30 |
| | | | | 0.737 | 0.280 | 5.76 | 4.23 | 0.000 | -20 | -2 | 28 |
| | | | | 0.850 | 0.288 | 5.52 | 4.12 | 0.000 | -18 | 6 | 22 |
| 0.019 | 0.004 | 127 | 0.001 | 0.842 | 0.288 | 5.54 | 4.13 | 0.000 | -26 | 24 | 60 |
| | | | | 0.995 | 0.520 | 4.85 | 3.79 | 0.000 | -26 | 12 | 64 |
| | | | | 1.000 | 0.852 | 3.89 | 3.24 | 0.001 | -24 | 4 | 66 |
| 0.007 | 0.002 | 153 | 0.000 | 0.961 | 0.391 | 5.16 | 3.95 | 0.000 | -22 | -86 | 38 |
| | | | | 1.000 | 0.599 | 4.56 | 3.63 | 0.000 | -20 | -72 | 38 |
| | | | | 1.000 | 0.662 | 4.21 | 3.44 | 0.000 | -22 | -76 | 48 |
| 0.000 | 0.000 | 242 | 0.000 | 0.973 | 0.415 | 5.09 | 3.92 | 0.000 | 28 | -78 | 44 |
| | | | | 1.000 | 0.662 | 4.31 | 3.49 | 0.000 | 26 | -90 | 34 |
| | | | | 1.000 | 0.709 | 4.08 | 3.36 | 0.000 | 34 | -86 | 36 |
| 0.009 | 0.002 | 148 | 0.000 | 0.981 | 0.423 | 5.03 | 3.88 | 0.000 | 52 | 10 | -32 |
| | | | | 1.000 | 0.633 | 4.49 | 3.59 | 0.000 | 46 | 10 | -40 |
| 0.017 | 0.004 | 131 | 0.001 | 1.000 | 0.599 | 4.59 | 3.65 | 0.000 | 46 | -62 | -12 |
| | | | | 1.000 | 0.601 | 4.54 | 3.63 | 0.000 | 44 | -60 | -4 |
| | | | | 1.000 | 0.662 | 4.26 | 3.47 | 0.000 | 38 | -60 | -10 |
| 0.000 | 0.000 | 236 | 0.000 | 1.000 | 0.662 | 4.36 | 3.52 | 0.000 | 42 | -22 | 48 |
| | | | | 1.000 | 0.662 | 4.29 | 3.48 | 0.000 | 34 | -28 | 66 |
| | | | | 1.000 | 0.662 | 4.26 | 3.47 | 0.000 | 34 | -26 | 58 |

**Table S6**. Second-level GLM results for the [Positive FC target > Baseline] contrast.

| cluster-level | | | | peak-level | | | | | coordinates | | |
|---|---|---|---|---|---|---|---|---|---|---|---|
| $p_{FWE\text{-}corr}$ | $q_{FDR\text{-}corr}$ | $k_E$ | $p_{uncorr}$ | $p_{FWE\text{-}corr}$ | $q_{FDR\text{-}corr}$ | T | ($Z_E$) | $p_{uncorr}$ | X | Y | Z |
| **Positive FC target > Baseline** | | | | | | | | | | | |
| 0.000 | 0.000 | 6907 | 0.000 | 0.001 | 0.017 | 10.91 | 5.77 | 0.000 | 40 | -14 | -2 |
| | | | | 0.008 | 0.024 | 9.34 | 5.39 | 0.000 | 62 | -22 | 20 |
| | | | | 0.012 | 0.024 | 9.08 | 5.32 | 0.000 | 62 | -14 | 16 |
| 0.000 | 0.000 | 3688 | 0.000 | 0.015 | 0.024 | 8.96 | 5.29 | 0.000 | -42 | -10 | -6 |
| | | | | 0.110 | 0.048 | 7.69 | 4.91 | 0.000 | -40 | -4 | -14 |
| | | | | 0.182 | 0.055 | 7.39 | 4.81 | 0.000 | -40 | -16 | 18 |
| 0.000 | 0.000 | 1260 | 0.000 | 0.111 | 0.048 | 7.68 | 4.90 | 0.000 | -0 | 12 | 36 |
| | | | | 0.545 | 0.119 | 6.38 | 4.44 | 0.000 | 4 | 2 | 40 |
| | | | | 0.921 | 0.205 | 5.54 | 4.08 | 0.000 | -14 | 10 | 36 |
| 0.008 | 0.002 | 135 | 0.000 | 0.138 | 0.053 | 7.55 | 4.86 | 0.000 | -24 | -66 | -22 |
| | | | | 0.712 | 0.152 | 6.05 | 4.30 | 0.000 | -20 | -62 | -30 |
| | | | | 0.994 | 0.300 | 5.06 | 3.85 | 0.000 | -12 | -72 | -18 |
| 0.010 | 0.002 | 130 | 0.000 | 0.216 | 0.058 | 7.25 | 4.76 | 0.000 | 14 | -68 | -22 |
| | | | | 0.772 | 0.153 | 5.92 | 4.25 | 0.000 | 12 | -74 | -16 |
| | | | | 0.929 | 0.205 | 5.51 | 4.07 | 0.000 | 8 | -64 | -16 |
| 0.000 | 0.000 | 458 | 0.000 | 0.331 | 0.083 | 6.87 | 4.62 | 0.000 | 30 | -60 | -26 |
| | | | | 0.751 | 0.152 | 5.97 | 4.27 | 0.000 | 40 | -54 | -30 |
| | | | | 0.993 | 0.300 | 5.06 | 3.86 | 0.000 | 44 | -54 | -38 |
| 0.004 | 0.001 | 154 | 0.000 | 0.758 | 0.152 | 5.95 | 4.26 | 0.000 | 48 | -4 | 52 |
| | | | | 1.000 | 0.697 | 4.05 | 3.31 | 0.000 | 52 | -2 | 44 |
| | | | | 1.000 | 0.769 | 3.96 | 3.26 | 0.001 | 36 | -6 | 48 |
| 0.017 | 0.004 | 116 | 0.001 | 0.813 | 0.163 | 5.83 | 4.21 | 0.000 | 8 | -38 | -30 |
| | | | | 0.998 | 0.326 | 4.90 | 3.78 | 0.000 | 4 | -28 | -22 |
| | | | | 1.000 | 0.550 | 4.32 | 3.47 | 0.000 | 12 | -46 | -34 |

**Table S7**. Second-level GLM results for the [Positive FC target < Baseline] contrast.

| cluster-level | | | | peak-level | | | | | coordinates | | |
|---|---|---|---|---|---|---|---|---|---|---|---|
| $p_{FWE-corr}$ | $q_{FDR-corr}$ | $k_E$ | $p_{uncorr}$ | $p_{FWE-corr}$ | $q_{FDR-corr}$ | T | ($Z_E$) | $p_{uncorr}$ | X | Y | Z |
| **Positive FC target < Baseline** | | | | | | | | | | | |
| 0.000 | 0.000 | 274 | 0.000 | 0.057 | 0.232 | 8.09 | 5.03 | 0.000 | -14 | -66 | 60 |
| | | | | 0.103 | 0.232 | 7.72 | 4.92 | 0.000 | -22 | -58 | 56 |
| | | | | 0.826 | 0.460 | 5.80 | 4.20 | 0.000 | -16 | -68 | 68 |
| 0.000 | 0.000 | 492 | 0.000 | 0.095 | 0.232 | 7.77 | 4.93 | 0.000 | -38 | -4 | 26 |
| | | | | 0.278 | 0.260 | 7.03 | 4.68 | 0.000 | -30 | -0 | 30 |
| | | | | 0.646 | 0.460 | 6.18 | 4.36 | 0.000 | -24 | -10 | 34 |
| 0.000 | 0.000 | 437 | 0.000 | 0.210 | 0.260 | 7.27 | 4.77 | 0.000 | 8 | -38 | 74 |
| | | | | 0.753 | 0.460 | 5.96 | 4.27 | 0.000 | 4 | -40 | 64 |
| | | | | 0.997 | 0.516 | 4.98 | 3.81 | 0.000 | -4 | -32 | 72 |
| 0.000 | 0.000 | 1374 | 0.000 | 0.439 | 0.341 | 6.60 | 4.52 | 0.000 | 16 | -60 | 68 |
| | | | | 0.739 | 0.460 | 5.99 | 4.28 | 0.000 | 22 | -72 | 52 |
| | | | | 0.807 | 0.460 | 5.85 | 4.22 | 0.000 | 24 | -62 | 52 |
| 0.000 | 0.000 | 222 | 0.000 | 0.798 | 0.460 | 5.87 | 4.23 | 0.000 | 42 | -28 | 36 |
| | | | | 0.984 | 0.468 | 5.20 | 3.92 | 0.000 | 46 | -28 | 54 |
| | | | | 1.000 | 0.664 | 4.54 | 3.59 | 0.000 | 54 | -18 | 50 |
| 0.010 | 0.003 | 128 | 0.000 | 0.906 | 0.460 | 5.59 | 4.10 | 0.000 | 24 | -70 | -8 |
| | | | | 0.977 | 0.462 | 5.26 | 3.95 | 0.000 | 32 | -70 | -6 |
| | | | | 0.981 | 0.462 | 5.23 | 3.93 | 0.000 | 26 | -78 | -12 |
| 0.001 | 0.000 | 198 | 0.000 | 0.939 | 0.460 | 5.47 | 4.05 | 0.000 | 18 | 22 | 42 |
| | | | | 0.990 | 0.468 | 5.12 | 3.88 | 0.000 | 26 | 22 | 56 |
| | | | | 0.997 | 0.525 | 4.94 | 3.80 | 0.000 | 24 | 8 | 44 |
| 0.023 | 0.006 | 110 | 0.001 | 0.947 | 0.460 | 5.44 | 4.04 | 0.000 | -4 | 24 | -0 |
| | | | | 0.997 | 0.516 | 4.97 | 3.81 | 0.000 | 6 | 18 | 4 |
| | | | | 1.000 | 0.600 | 4.70 | 3.67 | 0.000 | -6 | 18 | 8 |
| 0.001 | 0.000 | 205 | 0.000 | 0.952 | 0.460 | 5.42 | 4.02 | 0.000 | -44 | 34 | 38 |
| | | | | 0.980 | 0.462 | 5.24 | 3.94 | 0.000 | -40 | 22 | 34 |
| | | | | 0.999 | 0.564 | 4.85 | 3.75 | 0.000 | -34 | 14 | 38 |
| 0.002 | 0.001 | 170 | 0.000 | 0.982 | 0.462 | 5.22 | 3.93 | 0.000 | 54 | -10 | -12 |
| | | | | 0.992 | 0.468 | 5.09 | 3.87 | 0.000 | 62 | -14 | -12 |
| | | | | 0.992 | 0.468 | 5.09 | 3.87 | 0.000 | 54 | -32 | -18 |
| 0.002 | 0.001 | 174 | 0.000 | 0.993 | 0.468 | 5.08 | 3.86 | 0.000 | 22 | -8 | 34 |
| | | | | 0.998 | 0.525 | 4.94 | 3.79 | 0.000 | 24 | 2 | 24 |
| | | | | 1.000 | 0.674 | 4.51 | 3.57 | 0.000 | 16 | -8 | 26 |
| 0.024 | 0.006 | 109 | 0.001 | 1.000 | 0.647 | 4.58 | 3.61 | 0.000 | 46 | -64 | -8 |

**Table S8**. Second-level GLM results for the [ Positive FC control target > Baseline] contrast.

| cluster-level | | | | peak-level | | | | | coordinates | | |
|---|---|---|---|---|---|---|---|---|---|---|---|
| $p_{FWE-corr}$ | $q_{FDR-corr}$ | $k_E$ | $p_{uncorr}$ | $p_{FWE-corr}$ | $q_{FDR-corr}$ | T | $(Z_E)$ | $p_{uncorr}$ | X | Y | Z |
| **Positive FC *control* target > Baseline** | | | | | | | | | | | |
| 0.000 | 0.000 | 4361 | 0.000 | 0.002 | 0.034 | 10.72 | 5.61 | 0.000 | -46 | -14 | 20 |
| | | | | 0.012 | 0.062 | 9.45 | 5.32 | 0.000 | -30 | -30 | 14 |
| | | | | 0.103 | 0.094 | 7.99 | 4.92 | 0.000 | -50 | -30 | 10 |
| 0.000 | 0.000 | 4285 | 0.000 | 0.069 | 0.094 | 8.26 | 5.00 | 0.000 | 36 | -26 | 16 |
| | | | | 0.403 | 0.110 | 7.04 | 4.61 | 0.000 | 54 | -32 | 18 |
| | | | | 0.432 | 0.110 | 6.97 | 4.59 | 0.000 | 16 | -16 | 2 |
| 0.000 | 0.000 | 416 | 0.000 | 0.727 | 0.151 | 6.32 | 4.35 | 0.000 | -0 | 18 | 36 |
| | | | | 0.985 | 0.231 | 5.43 | 3.98 | 0.000 | -2 | -2 | 44 |
| | | | | 0.987 | 0.235 | 5.41 | 3.97 | 0.000 | -8 | 8 | 38 |
| 0.045 | 0.014 | 83 | 0.001 | 0.897 | 0.183 | 5.89 | 4.18 | 0.000 | -46 | -10 | 52 |
| | | | | 1.000 | 0.738 | 4.09 | 3.30 | 0.000 | -54 | -12 | 54 |

**Table S9**. Second-level GLM results for the [ Positive FC control target < Baseline] contrast.

| cluster-level | | | | peak-level | | | | | coordinates | | |
|---|---|---|---|---|---|---|---|---|---|---|---|
| $p_{FWE-corr}$ | $q_{FDR-corr}$ | $k_E$ | $p_{uncorr}$ | $p_{FWE-corr}$ | $q_{FDR-corr}$ | T | ($Z_E$) | $p_{uncorr}$ | X | Y | Z |
| **Positive FC *control* target < Baseline** | | | | | | | | | | | |
| 0.000 | 0.000 | 561 | 0.000 | 0.225 | 0.417 | 7.50 | 4.76 | 0.000 | -2 | -22 | 76 |
| | | | | 0.307 | 0.417 | 7.30 | 4.70 | 0.000 | -8 | -42 | 72 |
| | | | | 0.521 | 0.547 | 6.76 | 4.51 | 0.000 | 14 | -46 | 76 |
| 0.000 | 0.000 | 275 | 0.000 | 0.268 | 0.417 | 7.39 | 4.73 | 0.000 | 34 | -26 | 34 |
| | | | | 0.638 | 0.547 | 6.51 | 4.42 | 0.000 | 30 | -46 | 24 |
| | | | | 0.972 | 0.630 | 5.56 | 4.03 | 0.000 | 44 | -22 | 40 |
| 0.008 | 0.003 | 118 | 0.000 | 0.540 | 0.547 | 6.72 | 4.50 | 0.000 | 42 | 30 | -12 |
| | | | | 1.000 | 0.787 | 4.73 | 3.64 | 0.000 | 44 | 36 | -4 |
| | | | | 1.000 | 0.859 | 4.32 | 3.43 | 0.000 | 46 | 20 | -20 |
| 0.000 | 0.000 | 205 | 0.000 | 0.668 | 0.547 | 6.44 | 4.40 | 0.000 | -62 | -22 | -8 |
| | | | | 0.999 | 0.736 | 5.07 | 3.81 | 0.000 | -56 | -34 | -0 |
| | | | | 1.000 | 0.776 | 4.81 | 3.68 | 0.000 | -58 | -12 | -10 |
| 0.000 | 0.000 | 542 | 0.000 | 0.790 | 0.630 | 6.18 | 4.29 | 0.000 | 14 | 60 | 10 |
| | | | | 0.977 | 0.630 | 5.51 | 4.01 | 0.000 | -4 | 64 | 18 |
| | | | | 0.981 | 0.630 | 5.48 | 4.00 | 0.000 | 12 | 54 | 16 |
| 0.001 | 0.001 | 158 | 0.000 | 0.966 | 0.630 | 5.59 | 4.05 | 0.000 | 58 | 6 | -32 |
| | | | | 1.000 | 0.859 | 4.20 | 3.36 | 0.000 | 58 | -2 | -26 |
| | | | | 1.000 | 0.859 | 4.19 | 3.36 | 0.000 | 46 | 20 | -30 |

**Table S10**. Second-level GLM results for the [ Negative FC target < Negative FC control target] contrast.

| cluster-level | | | | peak-level | | | | | coordinates | | |
|---|---|---|---|---|---|---|---|---|---|---|---|
| $p_{FWE\text{-}corr}$ | $q_{FDR\text{-}corr}$ | $k_E$ | $p_{uncorr}$ | $p_{FWE\text{-}corr}$ | $q_{FDR\text{-}corr}$ | T | $(Z_E)$ | $p_{uncorr}$ | X | Y | Z |
| **Negative FC target < Negative FC *control* target** | | | | | | | | | | | |
| 0.000 | 0.000 | 256 | 0.000 | 0.773 | 0.725 | 5.77 | 4.24 | 0.000 | -8 | 28 | -16 |
| | | | | 0.997 | 0.725 | 4.87 | 3.80 | 0.000 | 8 | 44 | -16 |
| | | | | 1.000 | 0.788 | 4.50 | 3.60 | 0.000 | 4 | 32 | -12 |
| 0.001 | 0.001 | 180 | 0.000 | 0.844 | 0.725 | 5.62 | 4.17 | 0.000 | -16 | 56 | 22 |
| | | | | 0.999 | 0.788 | 4.69 | 3.70 | 0.000 | -4 | 66 | 18 |
| | | | | 1.000 | 0.788 | 4.54 | 3.62 | 0.000 | -16 | 66 | 16 |
| 0.024 | 0.009 | 109 | 0.001 | 0.942 | 0.725 | 5.33 | 4.03 | 0.000 | 8 | 50 | 26 |
| | | | | 0.978 | 0.725 | 5.14 | 3.94 | 0.000 | -6 | 42 | 32 |
| | | | | 1.000 | 0.865 | 4.22 | 3.44 | 0.000 | 2 | 46 | 32 |

**Table S11**. Second-level GLM results for the [[Negative FC target > Negative FC control target] < [Positive FC target > Positive FC control target]] contrast.

| cluster-level | | | | peak-level | | | | | coordinates | | |
|---|---|---|---|---|---|---|---|---|---|---|---|
| $p_{FWE-corr}$ | $q_{FDR-corr}$ | $k_E$ | $p_{uncorr}$ | $p_{FWE-corr}$ | $q_{FDR-corr}$ | T | $(Z_E)$ | $p_{uncorr}$ | X | Y | Z |
| **[Negative FC target > Negative FC *control* target] < [Positive FC target > Positive FC *control* target]** | | | | | | | | | | | |
| 0.006 | 0.006 | 181 | 0.000 | 0.144 | 0.416 | 5.28 | 4.68 | 0.000 | 4 | 22 | -18 |
| | | | | 0.572 | 0.524 | 4.69 | 4.24 | 0.000 | -8 | 26 | -18 |
| | | | | 0.838 | 0.524 | 4.40 | 4.02 | 0.000 | 4 | 16 | -12 |
| 0.000 | 0.000 | 528 | 0.000 | 0.483 | 0.524 | 4.78 | 4.31 | 0.000 | -4 | 66 | 18 |
| | | | | 0.928 | 0.524 | 4.26 | 3.91 | 0.000 | -14 | 44 | 6 |
| | | | | 0.968 | 0.524 | 4.16 | 3.83 | 0.000 | -12 | 54 | 18 |

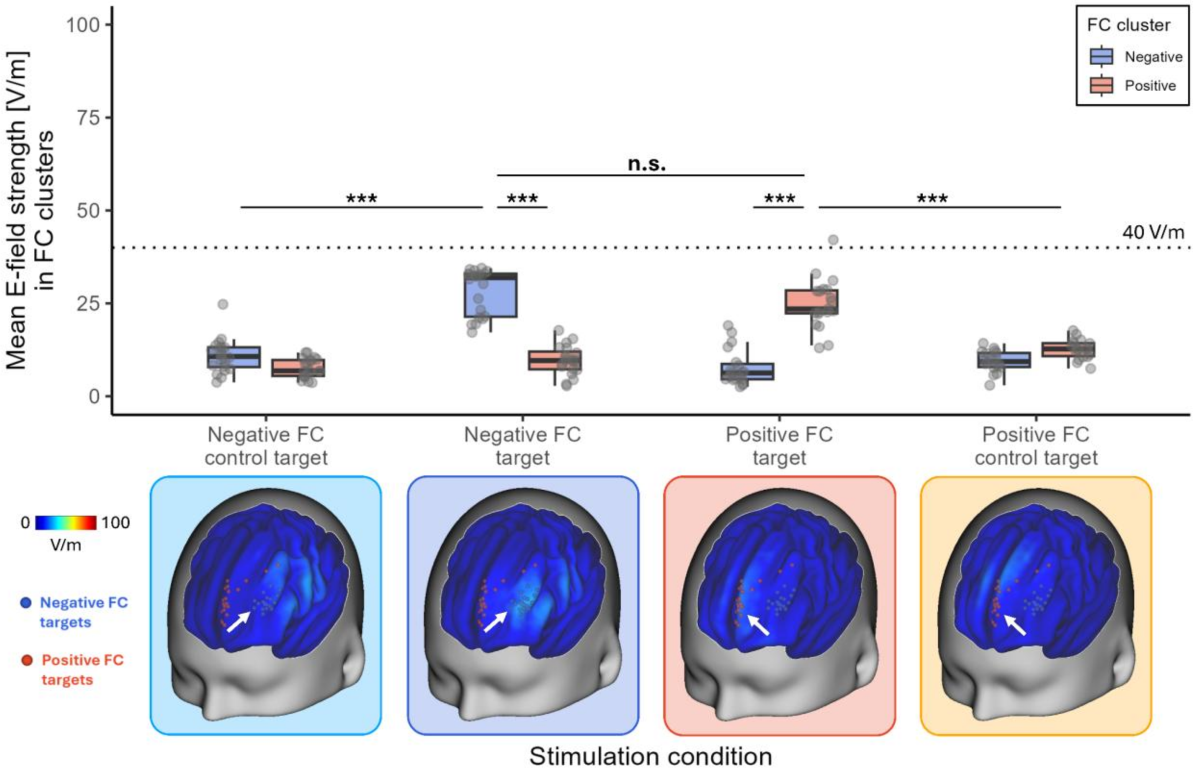


**Figure S1**. Average simulated TMS-induced electric fields per stimulation condition for low-intensity pre-pulses. Top: Boxplots of mean E-field values in the individually identified negative (blue) and positive (red) FC clusters (n = 20 for negative FC conditions, n = 19 for the positive FC target, n = 18 for the positive FC control target). The dotted line marks 40 V/m. Bottom: Sample-averaged simulated E-fields for each stimulation condition visualized on the cortical surface (same color scale as Figure 2C). Brackets indicate paired comparisons, Bonferroni-corrected for five comparisons: negative FC target vs. control (mean difference = 17.29 V/m [14.29, 20.30], $t(19) = 16.45$, $p_{Bonf} < .001$, $d = 3.05$ [1.85, 4.24]); negative vs. positive cluster during negative FC target stimulation (mean difference = 18.51 V/m [15.03, 21.98], $t(19) = 15.23$, $p_{Bonf} < .001$, $d = 3.41$ [1.83, 4.99]); negative vs. positive FC target (mean difference = 2.94 V/m [−0.87, 6.74], $t(18) = 2.22$, $p_{Bonf} = .196$, $d = 0.45$ [−0.13, 1.03]); positive vs. negative cluster during positive FC target stimulation (mean difference = 17.27 V/m [12.65, 21.90], Wilcoxon $V = 190$, $p_{Bonf} < .001$, $r = 1.00$); positive FC target vs. control (mean difference = 12.36 V/m [7.24, 17.47], $t(17) = 7.00$, $p_{Bonf} < .001$, $d = 2.38$ [0.57, 4.18]). ***$p_{Bonf} < .001$, n.s. = not significant ($p_{Bonf} > .05$).

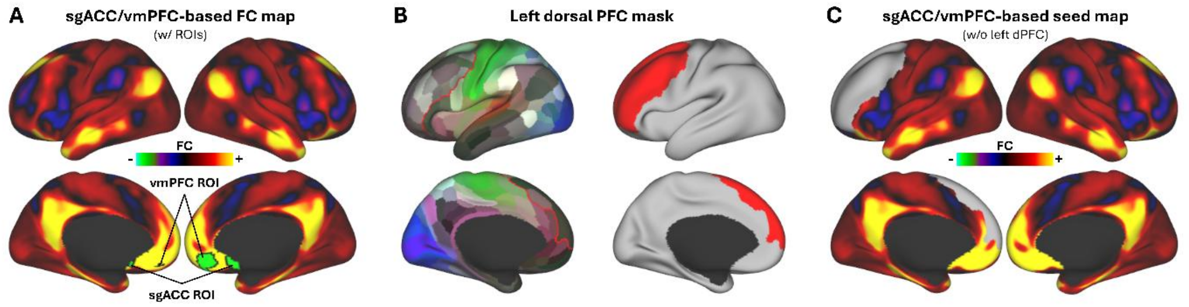


**Figure S2**. Seed map for individual FC analyses. **A**: Full normative FC map, based on a sample of n = 1003 participants from the Human Connectome Project (6), using sgACC (X = 5.9, Y = 16.3, Z = -9.8) (7) and vmPFC (X = 6, Y = 46, Z = -14) (8) regions of interest (10 mm radius) as seeds. **B**: Left dorsal PFC mask used as the search space area for TMS target clusters, made from the following parcels of the HCP atlas (51): L_8Ad_ROI, L_8Av_ROI, L_8BL_ROI, L_8BM_ROI, L_8C_ROI, L_9-46d_ROI, L_9a_ROI, L_9m_ROI, L_9p_ROI, L_10d_ROI, L_46_ROI, L_IFJa_ROI, L_IFJp_ROI, L_IFSa_ROI, L_IFSp_ROI, L_a9-46v_ROI, L_a10p_ROI, L_a47r_ROI, L_i6-8_ROI, L_p9-46v_ROI, L_p10p_ROI, L_p47r_ROI, L_s6-8_ROI. **C**: Final sgACC/vmPFC-based FC map from A, excluding the dorsal PFC area from B, used as seed map for individual FC analyses.

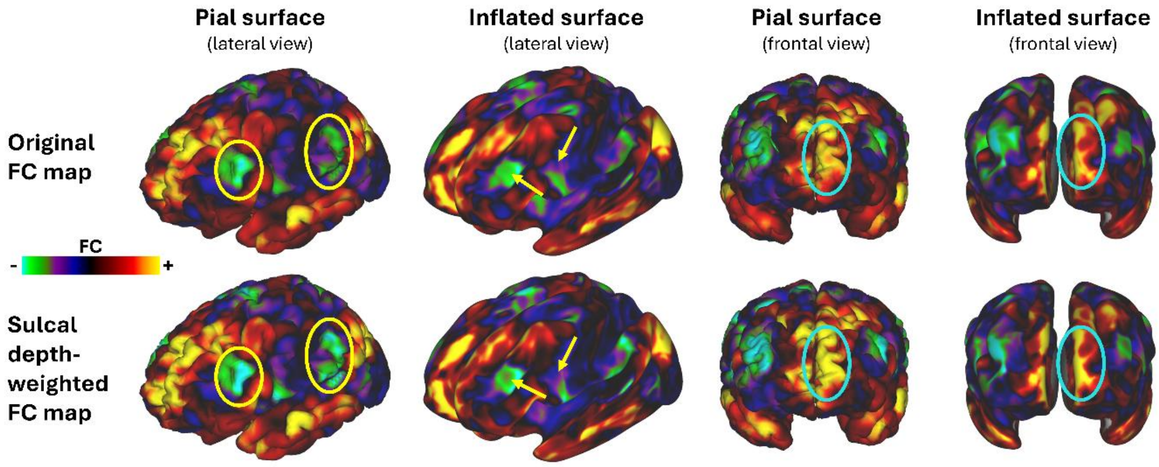


**Figure S3**. Illustration of sulcal-depth weighting (for one representative participant). Areas on the gyral crowns are upweighted, while areas within sulci receive a relatively lower weight. Compare areas marked by circles and arrows between upper (original FC map) and lower (sulcal depth-weighted FC map) row for subtle effects of sulcal-depth weighting.

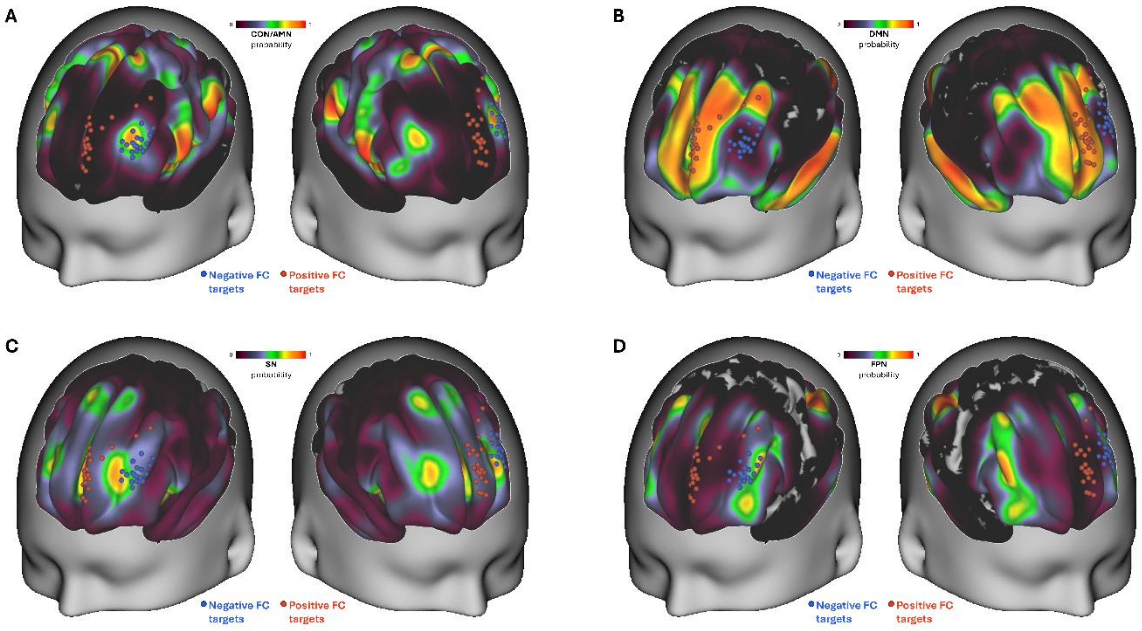


**Figure S4**. Distribution of the centers of negative (blue) and positive (red) FC target clusters relative to probability maps of (**A**) the cingulo-opercular/action mode network (CON/AMN), (**B**) the default mode network (DMN), (**C**) the salience network (SN), and (**D**) the frontoparietal network (FPN).

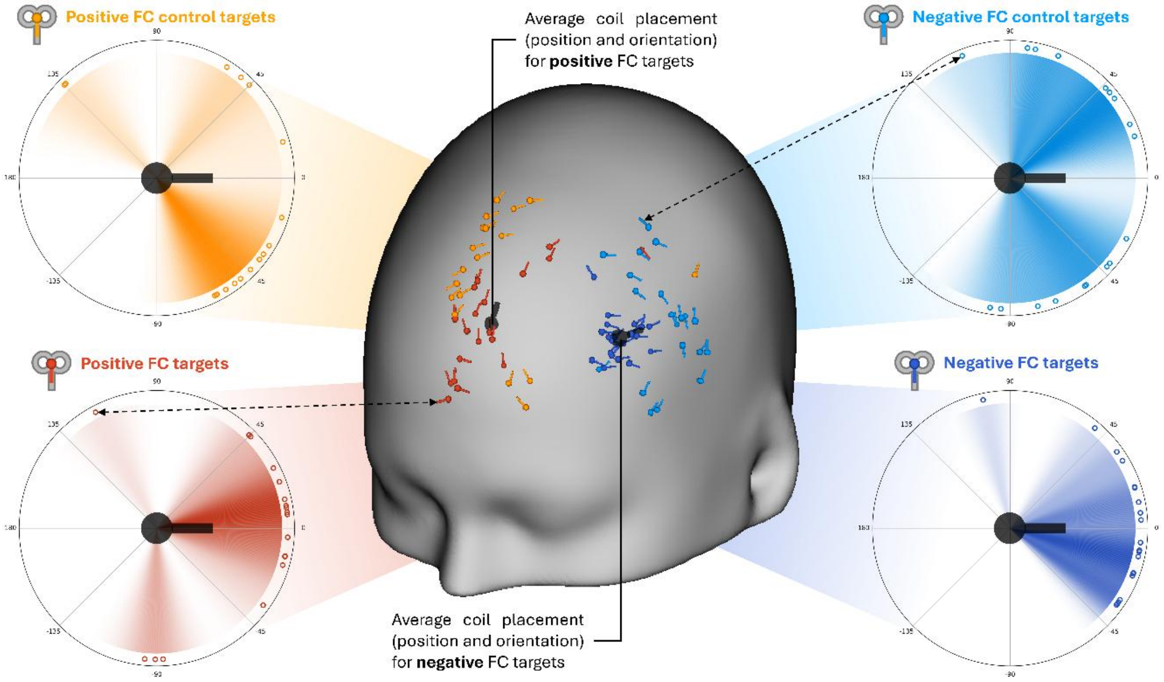


**Figure S5**. Interindividual variability of TMS coil angles/orientations for each of the four stimulation conditions: Negative FC targets (lower right), negative FC control targets (upper right), positive FC targets (lower left), positive FC control targets (upper left). Each polar plot indicates the angular deviation (in degrees) of individual TMS coil orientations (indicated by hollow dots at the edge of each polar plot) from the respective average coil orientation (larger black coil representations on the head surface and in the center of the polar plots) for negative FC targets (left) and positive FC targets (right). Dashed double arrows indicate corresponding coil placements on the head surface and polar plot.

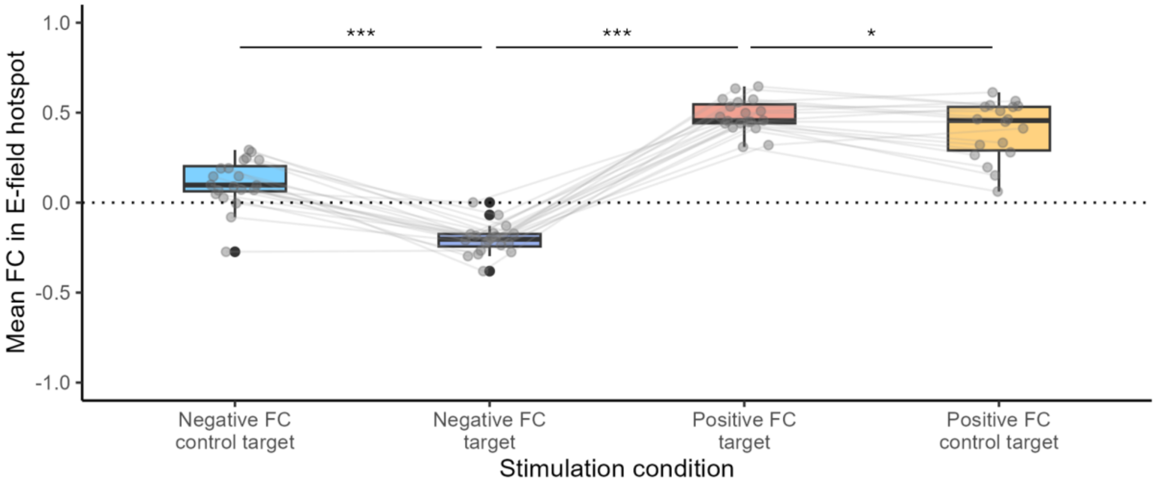


**Figure S6**. Mean functional connectivity (FC) values within condition-specific E-field hotspot areas. Note that, due to the generally unknown effective E-field threshold, each dot/value represents the average FC value across ten percentile-based E-field hotspot thresholds, from 99.0% to 99.9%, in steps of 0.1%. Lines connect values from the same participant ($n$ = 20 for negative FC conditions, $n$ = 19 for the positive FC target, $n$ = 18 for the positive FC control target). Brackets indicate paired t-tests: negative FC target vs. control ($t(19) = -9.85$, $p_{\text{Bonf}} < .001$, $d = -2.78$ [−4.34, −1.22]); negative vs. positive FC target ($t(18) = -22.79$, $p_{\text{Bonf}} < .001$, $d = -7.87$ [−12.77, −2.97]); positive FC target vs. control ($t(17) = 2.98$, $p_{\text{Bonf}} = .025$, $d = 0.56$ [0.05, 1.06]). ***$p_{\text{Bonf}} < .001$, *$p_{\text{Bonf}} < .05$.
